# The EGY1-SGR1 module controls chloroplast development and senescence by modulating photosynthetic functions

**DOI:** 10.1101/2025.11.10.687641

**Authors:** Alexey Shapiguzov, Andrea Trotta, Ilaria Mancini, Umama Hani, Nasrin Sultana, Triin Vahisalu, Cezary Waszczak, Eva-Mari Aro, Mikael Brosché

**Affiliations:** Organismal and Evolutionary Biology Research Program, Faculty of Biological and Environmental Sciences, Viikki Plant Science Centre, University of Helsinki, 00014 Helsinki, Finland; Natural Resources Institute Finland (Luke), Production Systems, 21500 Piikkiö, Finland; Molecular Plant Biology, Department of Life Technologies, University of Turku, Turku, Finland

## Abstract

The chloroplast plays essential roles not only in production of biochemical energy and metabolism, but also in plant development, including senescence. We describe a mutant with an unusual progression of senescence and identify it as an allele of *ethylene-dependent gravitropism-deficient and yellow-green 1* (*egy1-5*). *EGY1* encodes a chloroplast-localized metalloprotease that regulates chlorophyll metabolism. The single *egy1-5* mutant exhibits defects in various aspects of chloroplast function, including abnormal accumulation of photosynthetic complexes, defects in photosynthetic electron transfer chain and altered responses to the chemicals methyl viologen and lincomycin. Additionally, *egy1* lacks guard cell chloroplasts. We performed a suppressor mutant screen in the *egy1* background to identify new regulators of senescence and chloroplast development. We found two suppressors that restored green leaves and mapping revealed that both suppressors had mutations in *STAY-GREEN 1 (SGR1)*. *SGR1* encodes the Mg^2+^ dechelatase enzyme, which catalyses the first step in chlorophyll breakdown. Several defects in the *egy1* mutant were either fully or partially suppressed in *egy1 sgr1*. This enabled the function of EGY1 to be isolated, independent of the pleiotropic effects of chlorophyll degradation. Our findings suggest that chlorophyll homeostasis regulates the overall development and function of the chloroplast as well as plant senescence.

**Highlight:** Identifying new mutants with defects in chlorophyll metabolism and chloroplast development shows that proper chlorophyll homeostasis is necessary for maintaining functional chloroplasts.

## Introduction

In addition to photosynthetic light reactions and CO_2_ fixation, plant chloroplasts perform many essential functions, including the biosynthesis of amino acids, fatty acids, hormones, and secondary metabolites. Therefore, for optimal plant performance, the chloroplast requires co-ordination with other cell compartments. This involves receiving environmental and developmental signals. Conversely, the chloroplast is a key organelle in converting information about the light environment into molecular signals for other parts of the cell. Retrograde signalling from the chloroplast to the nucleus is required to synchronize transcriptional responses and assembly of photosynthetic protein complexes with stress and developmental responses, as well as energy metabolism (Gao *et al*., 2023; Richter *et al*., 2023). Via these mechanisms, the chloroplast develops in synchrony with plant’s life stages, throughout development from germination to senescence.

Chlorophyll is the main pigment responsible for capturing light, and an integral component of light-harvesting complexes and photosystems. Accumulation of free chlorophyll or its breakdown products should be avoided, as this can lead to the uncontrolled transfer of light energy to other molecules, generating for example reactive oxygen species. Therefore, the biosynthesis, breakdown and incorporation of chlorophyll into photosynthetic protein complexes must be tightly regulated (Wang and Grimm, 2015, 2021). It has also been suggested that chlorophyll itself regulates the stability of photosystem II (PSII) (Shimoda *et al*., 2016; Wang and Grimm, 2021).

Development of the chloroplast and assembly of photosynthetic protein complexes rely on nearly 20 proteases localized to the chloroplast (Nishimura *et al*., 2017). Their requirement for chloroplast development is illustrated, for example, by a variegated leaf phenotype of mutants deficient in the FTSH2 ATP-dependent metalloprotease (Nishimura *et al*., 2016). Another protease required for chloroplast development is a member of the S2P family of proteases, ETHYLENE-DEPENDENT GRAVITROPISM-DEFICIENT AND YELLOW-GREEN1 (EGY1) (Chen *et al*., 2005). The *egy1* mutant phenotypes include incorrect assembly of photosystem I (PSI) and deficient stabilization of PSII during light stress (Adamiec *et al*., 2021; Chen *et al*., 2021a; Qi *et al*., 2020; Sanjaya *et al*., 2021a; Sanjaya *et al*., 2021b).

Leaf senescence includes controlled degradation of chloroplasts and chlorophyll for recycling of nitrogen and nutrients. *STAY-GREEN 1* (*SGR1*, formerly called *NON-YELLOWING 1, NYE1*), encodes a Mg^2+^ dechelatase, that catalyses the first step of chlorophyll *a* breakdown during senescence (Shimoda *et al*., 2016). In the hormonal regulation of senescence, several transcription factors from ethylene, jasmonic acid and abscisic acid pathways directly regulate expression of *SGR1* and other genes encoding chlorophyll catabolic enzymes (Woo *et al*., 2019). For example, EIN3 (ETHYLENE-INSENSITIVE3), a key transcription factor in ethylene signalling binds to the promoter of *SGR1* (Qiu *et al*., 2015), as do the jasmonic acid signalling transcription factors MYC2/MYC3/MYC4 (Zhu *et al*., 2015), and the abscisic acid signalling transcription factors ABF2/ABF3/ABF4 (Gao *et al*., 2016). Collectively, this suggests that plant hormones act as direct regulators of senescence by targeting the breakdown of chlorophyll.

BALANCE of CHLOROPHYLL METABOLISM1 (BCM1) is a scaffold protein that coordinates both biosynthesis and degradation of chlorophyll (Wang *et al*., 2020). In the chlorophyll degradation pathway, BCM1 interacts with EGY1, targeting SGR1 for proteolytic degradation and thereby controlling the rate of chlorophyll degradation (Fu *et al*., 2025).

Here we describe the isolation of a mutant with unusual premature senescence and defects in chlorophyll metabolism. Mapping of the mutant identified an allele of *egy1* (*egy1-5*). Using *egy1-5* in an unbiased suppressor mutant screen, we demonstrate that the regulatory pair *EGY1-SGR1* regulates not only chlorophyll metabolism but also the stability of PSII, energy signalling, retrograde signalling and chloroplast development.

## Results

### Identification of egy1-5

We isolated a spontaneous Arabidopsis mutant in Col-0 background, displaying pale green leaves and an unusual type of senescence, characterised by premature loss of chlorophyll and eventually almost complete chlorosis (Figures 1A, B). Identification of the causative mutation revealed a 4 base pair deletion in *EGY1* gene (Supplementary Fig. S1). We refer to this mutant allele as *egy1-5*. To confirm the correct identification of the mutation, we performed an allelism test with the T-DNA line *egy1-2*. The F1 plants displayed pale green and yellowing leaves similar to those of the parents (Figure 1C). Despite the striking chlorotic phenotype of mature plants, they were otherwise morphologically similar to Col-0 and successfully produced seeds.

**Figure 1.**
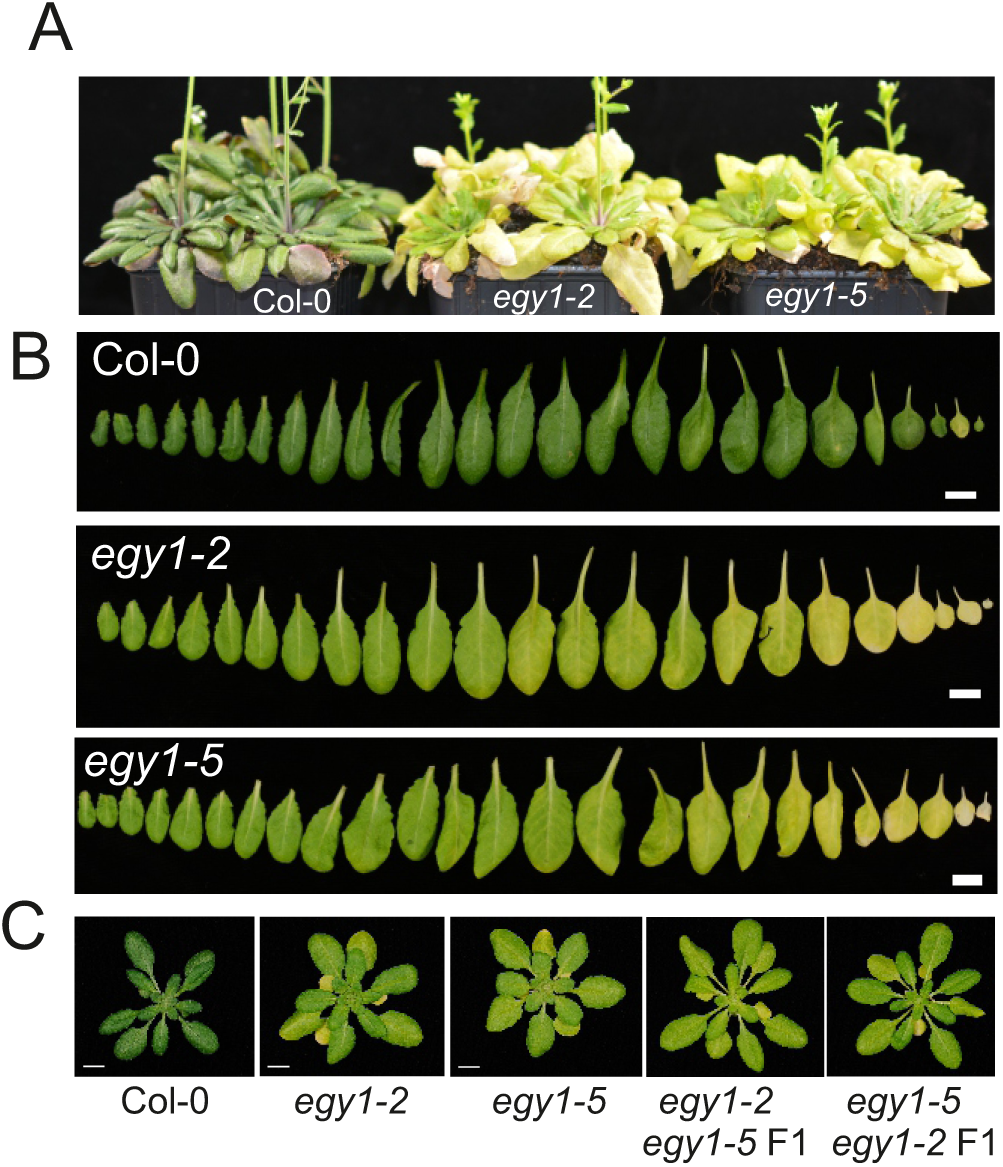
Identification of *egy1-5*. (A) Six-week-old Col-0, *egy1-2* and *egy1-5*. (B) Detached leaves from four-week-old plants. (C) Allelism test between *egy1-2* and *egy1-5*, showing four-week-old Col-0, *egy1-2*, *egy1-5*, *egy1-2 egy1-5* F1 and *egy1-5 egy1-2* F1 plants. Scale bar = 1 cm. See also Supplementary Fig. S1A for location of the four base pair deletion in *egy1-5*.

### Chlorosis of egy1 is rescued by the loss of function of STAYGREEN1

To identify new regulators of chloroplast development and chlorophyll metabolism, we conducted a suppressor mutant screen. Seeds of *egy1-5* were mutagenized with ethyl methane sulfonate (EMS) and M2 seeds were screened for plants with restored greener leaves. Two suppressors were selected for further characterization (Figure 2A). In both suppressor 1 (S1) and suppressor 2 (S2), the restored greener leaves became more apparent in older plants (Figure 2B). To identify the causative mutation of S2, we applied the SHOREmap backcross strategy (Hartwig *et al*., 2012). For this, S2 was backcrossed to *egy1-5*. The resulting BC1F1 had *egy1-5*-like phenotype indicating recessive inheritance. Analysis of SNP frequency in DNA sample obtained from bulked BC1F2 plants with restored green leaf colour revealed a high frequency (frequency = 1) G to A mutation at position 452 in *AT4G22920* encoding SGR1 (Supplementary Fig. S1B). Examination of the *SGR1* coding sequence in S1 revealed a G to A mutation at position 472. The new mutant alleles for *sgr1* were renamed *sgr1-3* (S1) and *sgr1-4* (S2). To confirm correct identification of the causative gene, we obtained another allele *sgr1-1* (also known as *nye1*, (Ren *et al*., 2007)), and crossed it with *egy1-5* (Figure 2A). The resulting double mutant *egy1-5 sgr1-1* also produced greener leaves in mature plants. We then complemented S1 and S2 with the wildtype (Col-0) *SGR1* gene, which restored the original pale and yellowing leaf phenotype (Figure 2B). Phylogenetic analysis of SGR from diverse plant species identified a conserved motif 1 (Jiao *et al*., 2020). The new identified SGR1 mutations lead to amino acids substitutions at this motif, R151Q and E158K (Fig 2B, inset). The arginine residue at position 151 was shown to be required for Mg^2+^ dechelatase activity of SGR1 (Xie *et al*., 2019).

**Figure 2.**
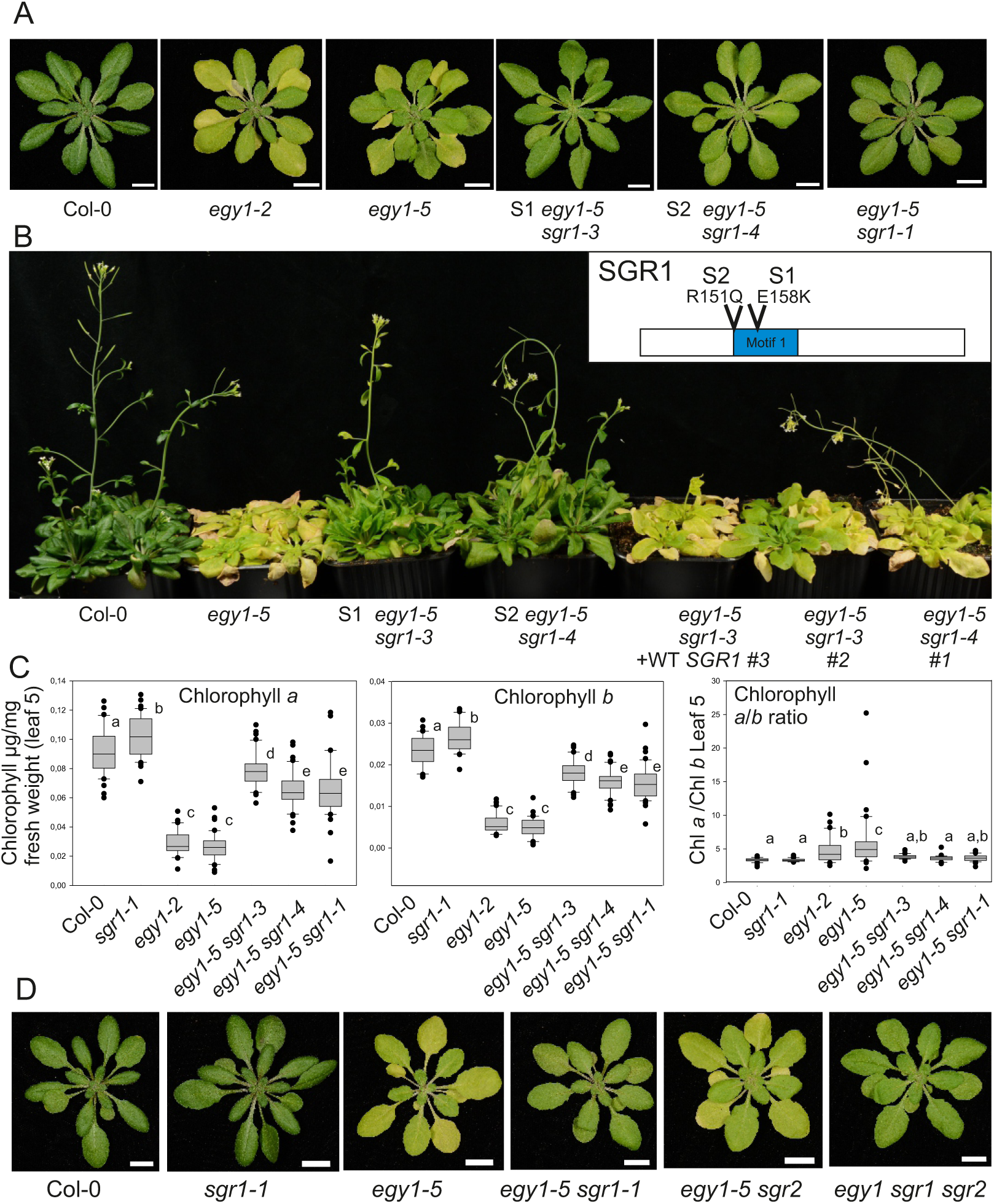
Identification of suppressor mutants that restore green phenotype in *egy1-5* background. (A) Four-week-old Col-0, *egy1-2*, *egy1-5*, *egy1-5 sgr1-3* (suppressor 1, S1), *egy1-5 sgr1-4* (suppressor 2, S2) and *egy1-5 sgr1-1*. (B) Genetic complementation in five-week-old plants. The *SGR1* gene including the native promoter was transformed into *egy1-5 sgr1-3* and *egy1-5 sgr1-4*. The inset shows the position of the mutations in the SGR1 amino acid sequence for *sgr1-3* (E158K) and *sgr1-4* (R151Q). (C) Chlorophyll was extracted from leaf 5 of four-week-old plants. The data shows chlorophyll *a* and *b*, and the *a*/*b* ratio from three biological repeats, N = 40 leaves. See also Supplementary Fig. S2 for chlorophyll measurements with Dualex. The box plots indicate the median, upper and lower quartile, and dots as outliers. Genotypes with different letters are statistically different (1-way ANOVA and Tukey’s multiple comparisons test). (D) Photos of four-week-old Col-0, *sgr1-1*, *egy1-5*, *egy1-5 sgr1-1*, *egy1-5 sgr2* and *egy1-5 sgr1-1 sgr2*. Scale bar = 1 cm.

As SGR1 catalyses the first step in chlorophyll degradation, we measured chlorophyll content *in vivo* and *in vitro* (Figure 2C, Supplementary Fig. S2). Both *egy1-2* and *egy1-5* had reduced levels of chlorophylls *a* and *b*, the decrease in chlorophyll *b* was more pronounced (Figure 2C), which is consistent with previous results (Adamiec *et al*., 2021). In the suppressor lines, chlorophyll levels and the chlorophyll *a*/*b* ratio were nearly fully restored.

In addition to SGR1, STAY-GREEN 2 (SGR2) also participates in chlorophyll breakdown (Sakuraba *et al*., 2014). To investigate possible genetic redundancy between SGR1 and SGR2, we generated the double mutant *egy1-5 sgr2* and the triple mutant *egy1-5 sgr1 sgr2*. The visual appearance of *egy1-5 sgr1 sgr2* was similar to *egy1-5 sgr1*, while *egy1-5 sgr2* was like *egy1-5* (Figure 2D). This suggests that SGR2 does not contribute to chlorophyll breakdown in the *egy1* mutant.

Next, we assessed whether other types of chloroplast defects associated with chlorosis could be rescued by *sgr1*. The mutant *var2* (*variegated2*), which lacks the thylakoid membrane protease FTSH2, exhibits defects in chloroplast development that lead to a variegated leaf phenotype (Nishimura *et al*., 2016). We generated *var2 sgr1* double mutant (Supplementary Fig. S3A). Unlike in the double *egy1 sgr1* mutant, loss of SGR1 function did not rescue the chlorotic phenotype of *var2*.

These experiments revealed the specific roles played by EGY1 and SGR1 in controlling plant senescence, establishing an experimental system in which leaf chlorophyll content can be modulated through genetic manipulations with *EGY1* and *SGR1*. Using this system, we studied the role of chlorophyll metabolism in various cellular processes, including photosynthetic electron transfer, the stability of photosynthetic protein complexes, responses to photosynthetic inhibitors, retrograde signalling, senescence and chloroplast development.

### Photosynthetic light reactions are impaired in *egy1* and partially restored in *egy1 sgr1*

SGR1 interacts directly with the light-harvesting complex II (LHCII) to control the breakdown of chlorophyll (Sakuraba *et al*., 2012), and the SGR1-mediated chlorophyll breakdown has been proposed as a regulator of PSII stability (Shimoda *et al*., 2016). Flash-induced chlorophyll fluorometry (OJIP kinetics) enables to probe the operation of the photosynthetic electron transfer chain. The OJIP kinetics of both *egy1-2* and *egy1-5* displayed lower relative contribution of O-J and J-I phases to overall fluorescence rise (Figure 3A). These phases correspond to the processes occurring in PSII (both the O-J and J-I rise) and in downstream electron acceptors of PSII (J-I rise) (Stirbet and Govindjee, 2011). For quantification, we extracted the relative fluorescence at 2 and 30 ms, which correspond to the O-J and J-I phases, accordingly (Figure 3B). The defects were partially restored to wildtype levels in the suppressor mutants *egy1 sgr1*. Importantly, *sgr1-1* single mutant had wildtype OJIP kinetics (Supplementary Fig. S4). This suggests alterations in PSII function and the redox state of the photosynthetic electron transfer chain in *egy1* that were partially reversed in *egy1 sgr1*.

**Figure 3.**
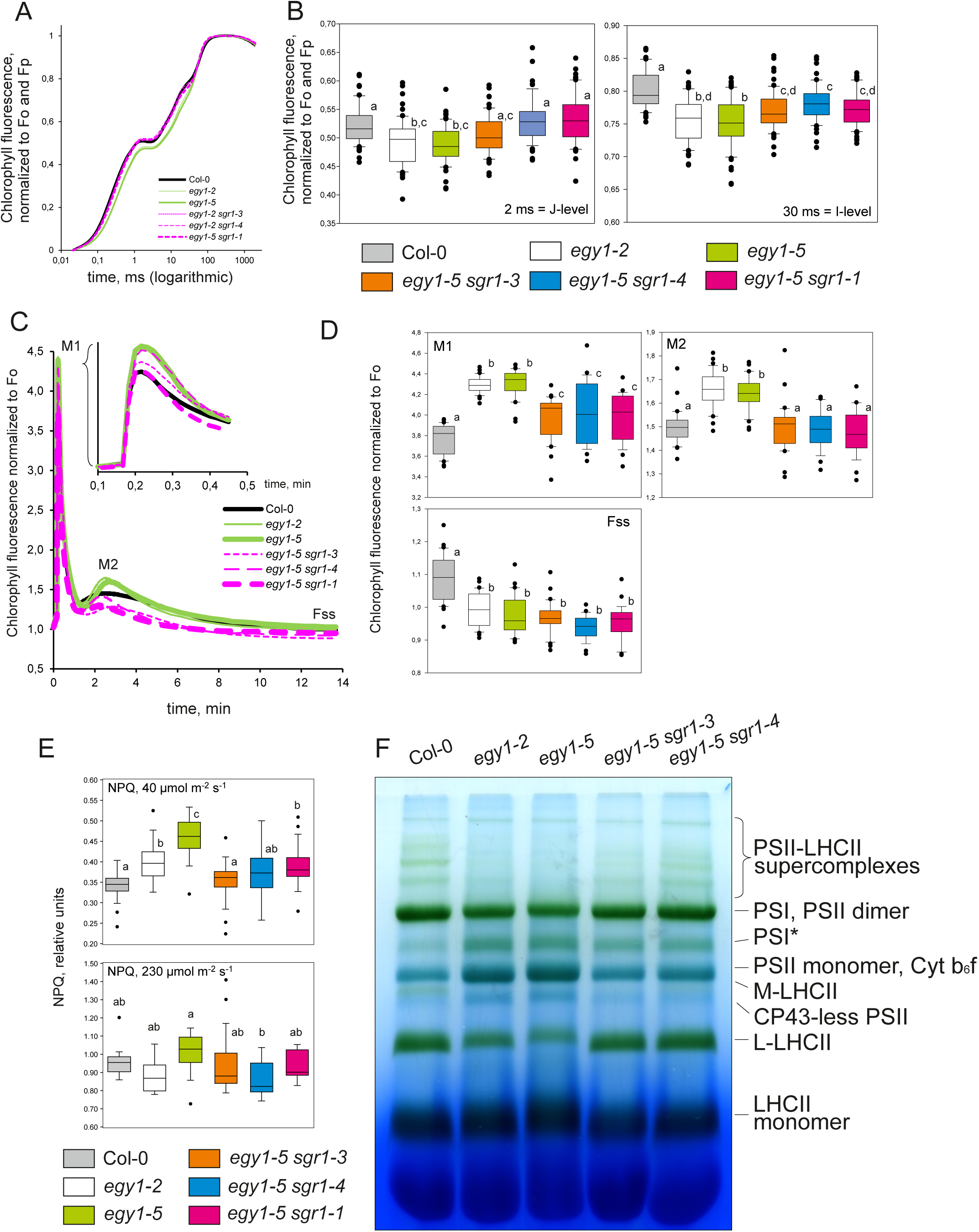
Photosynthetic electron transfer and composition of photosynthetic complexes in *egy1* and *egy1 sgr1*. (A) Kinetics of OJIP chlorophyll fluorescence rise, normalized to Fo and Fp. The curves represent the average of five plants. (B) OJIP data was extracted for the 2 and 30 ms time points from 12 biological repeats, N = 60 leaves. See also Supplementary Fig. S4 for OJIP data for the *sgr1* single mutant. (C) Chlorophyl fluorescence (Kautsky kinetics), from dark adapted plants exposed to low-intensity (80 µmol photons m^−2^ s^−1^) blue light (450 nm) inside the IMAGING-PAM M-Series. The curves represent one set of plants. The inset shows the early timescale of light treatment. (D) Several Kautsky kinetics parameters, M1, M2 and Fss, were quantified from six biological repeats, N = 27-32 plants. (E) NPQ was measured in 35 individual plants for actinic light 40 µmol m^−2^ s^−1^, and 15 individual plants for actinic light 230 µmol m^−2^ s^−1^. Letters indicate statistical significance (Tukey test, p < 0.05). The box plots indicate the median, upper and lower quartile, and dots as outliers. Genotypes with different letters are statistically different (1-way ANOVA and Tukey’s multiple comparisons test). (F) Separation by lpBN-PAGE of thylakoid protein complexes isolated from 28-day-old plants. The major protein complexes are indicated.

To explore acclimation of photosynthesis to changing light environment, we next exposed dark-acclimated plants to low-intensity light and measured the kinetics of chlorophyll fluorescence induction and relaxation (Kautsky kinetics). Oscillations in Kautsky kinetics are sensitive to various photosynthetic processes including light harvesting, photosynthetic electron flow and carbon assimilation (Stirbet and Govindjee, 2011; Stirbet *et al*., 2014; Tyystjärvi *et al*., 1999). We observed two distinct fluorescence maxima appearing at approximately 12 seconds (M1) and 2.5 minutes (M2) into the light treatment (Figure 3C). At both peaks, fluorescence was significantly higher in *egy1-2* and *egy1-5*, and this was partially (M1) or fully (M2) restored to wildtype levels in the suppressor mutants *egy1 sgr1* (Figure 3D).

Interestingly, steady-state fluorescence (Fss) was lower in *egy1-2* and *egy1-5* than in the wildtype, and it was not restored in *egy1 sgr1* (Figure 3D). This prompted us to assess non-photochemical quenching (NPQ) under two different light intensities (Figure 3E). In low light, NPQ was higher in *egy1,* and restored to wildtype levels in the suppressors. Overall, several chlorophyll fluorescence-based assays showed compromised functionality of photosynthetic apparatus in *egy1,* which was partially or fully restored in *egy1 sgr1*. This indicates a link between chlorophyll breakdown and the proper functioning of photosynthetic light reactions.

### Composition of photosynthetic complexes is affected in *egy1* and partially restored in *egy1 sgr1*

The distinct chlorophyll fluorescence phenotypes indicated problems with photosynthesis in *egy1-2* and *egy1-5*. To assess the composition of photosynthetic protein-pigment complexes, we isolated and solubilized thylakoids with β-dodecyl-n-maltoside (β-DM) and performed large-pore blue native gel electrophoresis (lpBN-PAGE). Both *egy1* mutants exhibited considerably reduced abundance of LHCII-containing complexes including PSII-LHCII supercomplexes and LHCII trimers (Figure 3F) (Chen *et al*., 2021a; Rantala *et al*., 2020). The defects worsened with plant age, becoming more pronounced at 35 days after germination (Supplementary Fig. S5) than after 28 days (Figure 3E). Thus, the absence of *EGY1* affected the abundance of LHCII and related protein complexes.

Next, we analysed the subunit composition of thylakoid protein complexes under denaturing conditions in the second dimension, stained the gels with SYPRO Ruby to detect total protein, or with Pro-Q Diamond to detect phosphoproteins (Supplementary Fig. S6). The analyses revealed similar protein composition in all the tested lines, including the individual core subunits of photosystems and the LHCA and LHCB subunits, although with lower accumulation of PSI, LHCII and LHCI in *egy1* mutants. Notably, the LHCB1 and LHCB2 proteins exhibited higher levels of phosphorylation in *egy1* mutants compared to the wildtype and the suppressor line *egy1-5 sgr1-4* (Supplementary Fig. S6). Phosphorylated LHCII is known to have higher affinity to PSI than PSII (Grieco *et al*., 2015). Lower accumulation of the PSI-LHCI-LHCII complex was observed in *egy1* mutants (Qi et al 2020), suggesting the existence of compensatory light-harvesting processes in *egy1*. Analysis of the protein composition of the L-LHCII region in lpBN-PAGE by mass spectrometry, including the detached LHCI belt (Grebe *et al*., 2019), revealed that no LHCII or LHCI isoforms were specifically missing in *egy1* or the suppressor line (Table 1, Supplementary Table S1). Furthermore, the L-LHCII region in *egy1* mutants was found to be present in complexes with an unusual subunit composition, including LHCB4, LHCB5 and LHCB6, which are not found in the L-LHCII band in Col-0 (Supplementary Fig. 6). Interestingly, the LHCB8 isoform (AT2G40100, also known as LHCB4.3 (Grebe *et al*., 2019)) was found in this band in mutant lines only.

**Table 1:** List of LHC proteins identified in L-LHCII bands separated by lpBN-PAGE.

| Gene symbol | Locus identifier | Coverage (%) | # peptides | #PSMs | #Unique peptides | Col-0<br>#Unique Peptides (#PSM) | *egy1-5*<br>#Unique Peptides (#PSM) | *egy1-5 sgr1-3*<br>#Unique Peptides (#PSM) |
| --- | --- | --- | --- | --- | --- | --- | --- | --- |
| LHCII isoforms |  |  |  |  |  |  |  |  |
| LHCB1.2 | AT1G29910 | 91 | 15 | 391 | 1 | 1(1) | 1(1) | 1(1) |
| LHCB1.3 | AT1G29930 | 90 | 15 | 391 | 1 | 1(1) | 1(1) | 1(1) |
| LHCB1.4 | AT2G34430 | 70 | 14 | 287 | 4 | 4(8) | 4(9) | 4(11) |
| LHCB1.5 | AT2G34420 | 77 | 14 | 381 | 1 | 1(1) | 1(1) | 1(2) |
| LHCB2* | AT3G27690<br>AT2G05100<br>AT2G05070 | 55 | 12 | 321 | 7 | 5(52) | 4(43) | 7(36) |
| LHCB3 | AT5G54270 | 69 | 13 | 85 | 12 | 11(32) | 5(17) | 10(26) |
| LHCB4.1 | AT5G01530 | 42 | 13 | 61 | 6 | 6(12) | 5(8) | 6(7) |
| LHCB4.2 | AT3G08940 | 50 | 17 | 66 | 10 | 9(12) | 9(11) | 9(10) |
| LHCB8 | AT2G40100 | 32 | 7 | 14 | 5 | - | 3(3) | 2(2) |
| LHCB5 | AT4G10340 | 62 | 18 | 69 | 18 | 18(28) | 16(25) | 13(16) |
| LHCB6 | AT1G15820 | 31 | 10 | 42 | 10 | 10(16) | 10(15) | 8(11) |
| LHCI isoforms |  |  |  |  |  |  |  |  |
| LHCA1 | AT3G54890 | 20 | 7 | 30 | 7 | 6(11) | 5(10) | 7(9) |
| LHCA2 | AT3G61470 | 10 | 1 | 5 | 1 | 1(2) | 1(1) | 1(2) |
| LHCA3 | AT1G61520 | 25 | 5 | 19 | 5 | 4(7) | 6(9) | 3(4) |
| LHCA4 | AT3G47470 | 36 | 7 | 29 | 7 | 5(7) | 7(12) | 6(10) |

\*For the three genes encoding LHCB2 isoforms, no unique peptides that could discriminate one from the other three proteins were identified, thus the unique peptides present only in the LHCB2 isoforms are equally valid for any of the three LHCB2 loci.

The wildtype composition of photosynthetic complexes was largely restored in *egy1 sgr1* (Figure 3F, Supplementary Fig. S5-S6). The levels of the LHCII complexes, which contain most of the cellular chlorophyll, returned to wildtype levels. However, the suppression was incomplete: the PSI assembly intermediate (PSI*) and PSII monomers were still more abundant in the suppressor lines than in the wildtype. Thus, the absence of EGY1 resulted in the severe depletion of the main chlorophyll-binding protein pigment complex, LHCII, and its larger assemblies, and this defect was partially rescued by the loss of SGR1.

### The contrasting reactions of *egy1* to photodamage are partially restored in *egy1 sgr1*

Various chemicals can be used to dissect specific photosynthetic functions. Methyl viologen (MV, also known as paraquat) catalyses electron transfer from PSI to molecular oxygen, leading to production of damaging reactive oxygen species. In a time series assay in which leaf discs were incubated in 0.25 µM MV (Shapiguzov and Kangasjarvi, 2022; Shapiguzov *et al*., 2019), *egy1-5* showed an extremely rapid drop in PSII maximal quantum efficiency (Fv/Fm), indicating sensitivity to damage to the photosynthetic electron transfer chain (Figure 4A). We quantified the data from the 4-hr time point (Figure 4B). In the suppressor mutant *egy1-5 sgr1-1,* the wildtype sensitivity to MV was partially restored compared to the *egy1-5* single mutant. This suggests that not only leaves are greener in *egy1-5 sgr1*, but also chloroplast reactive oxygen species metabolism is restored compared to single *egy1* mutant. In contrast, *sgr2* could not suppress the MV sensitivity of *egy1-5* (Figure 4B). Interestingly the extreme MV sensitivity of *var2*, which was visible even under low doses of MV, was partially restored in *var2 sgr1* (Supplementary Figs. S3B, C).

**Figure 4.**
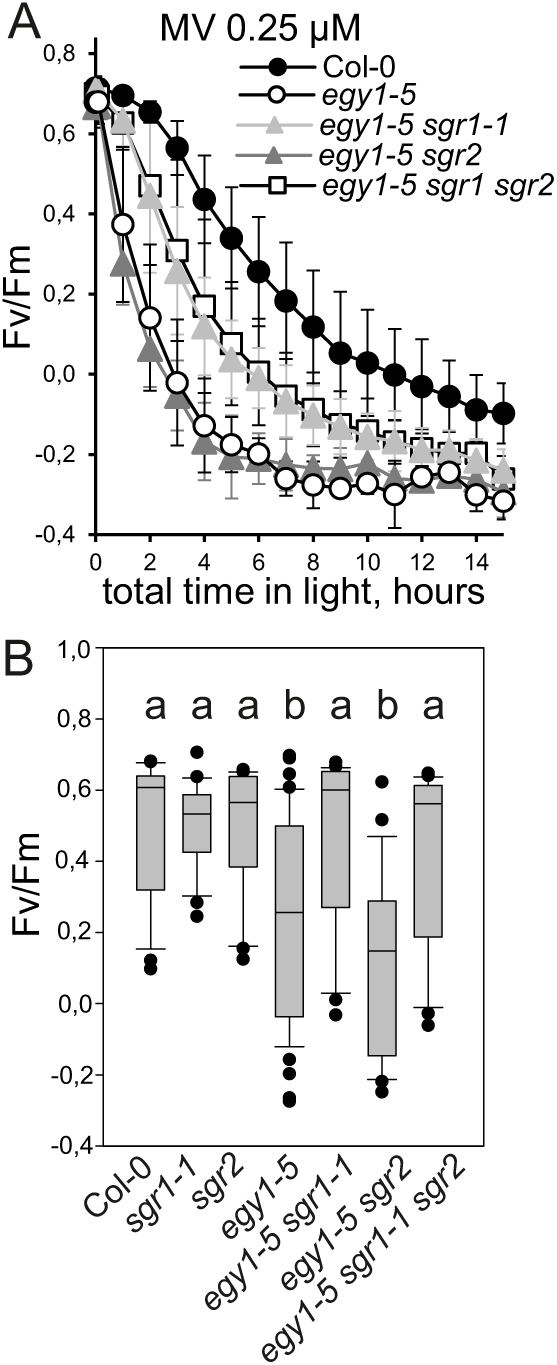
The response to MV. (A) Fv/Fm measured in leaf discs from four-week-old Col-0, *egy1-5*, *egy1-5 sgr1*, *egy1-5 sgr2, egy1-5 sgr1 sgr2, sgr1* and *sgr2* exposed to repeated cycles of blue light (448nm, 80 µmol m^−2^ s^−1^) and 0.25 µM MV. N = 5 leaf discs, the error bars indicate the SD. (B) Data from the 4-hr timepoint was quantified from five biological repeats, N=25 leaf discs. The box plot indicates the median, upper and lower quartile, and dots as outliers. Genotypes with different letters are statistically different (1-way ANOVA and Tukey’s multiple comparisons test).

Another chemical that has been used to investigate the function of the photosynthetic electron transfer is lincomycin, which inhibits translation in the chloroplast. The application of lincomycin prevents the *de novo* synthesis of chloroplast-encoded proteins, including the PSII D1 protein, which leads to impaired turnover of PSII (Tikkanen *et al*., 2014). We used the same time series assay with leaf discs as for MV, using 1 mM lincomycin (Supplementary Figs. S7A-D). We subsequently extracted data from the 4-hr timepoint (Supplementary Fig. S7E). As anticipated, lincomycin treatment led to decreased Fv/Fm in Col-0, but unexpectedly, *egy1-2* and *egy1-5* were resistant to the treatment. The increased tolerance of *egy1* to lincomycin may be linked to decreased light harvesting in these mutants. The fact that this trait coincides with greatly decreased resistance to MV, likely indicates complex changes in the maintenance of photosynthetic complexes that go beyond the scope of this study. Importantly, both phenotypes were partially rescued by *sgr1*.

### Impact of chlorophyll metabolism on retrograde signalling

As the chloroplast contains proteins encoded by both the nuclear and chloroplast genomes, the correct functioning of this organelle requires retrograde signalling from the chloroplast to the nucleus to coordinate the expression of photosynthesis-associated nuclear genes (PhANGs).

We tested genes related to PhANG, jasmonic acid, energy, and retrograde signalling in four-week-old plants using real-time quantitative PCR (qPCR). Modest changes in transcript levels were observed for most genes tested (Figure 5A, Supplementary Fig. S8). However, the *egy1-2* and *egy1-5* mutants displayed significantly higher transcript levels for senescence-related gene *DIN10* (*DARK INDUCIBLE 10*) (Fujiki *et al*., 2001)), jasmonic acid signalling gene *JAO3* (*JASMONATE-INDUCED OXYGENASE3*) (Caarls *et al*., 2017)), and sugar metabolism-related gene *KINBETA1* (a SnRK1 kinase subunit, which acts as a master regulator of plant metabolism (Li *et al*., 2009)). The transcript levels of *DIN10* serve as an indicator of the sugar status in plants (Fujiki *et al*., 2001). Increased transcript levels for both *DIN10* and *KINBETA1* suggested perturbed carbohydrate metabolism in *egy1*. Notably, the suppressor mutants exhibited complete restoration of *DIN10* and *JAO3* transcript levels to wildtype levels, suggesting that prevention of chlorophyll breakdown restores not only internal chloroplastic processes, but also molecular signalling pathways outside of the chloroplast.

**Figure 5.**
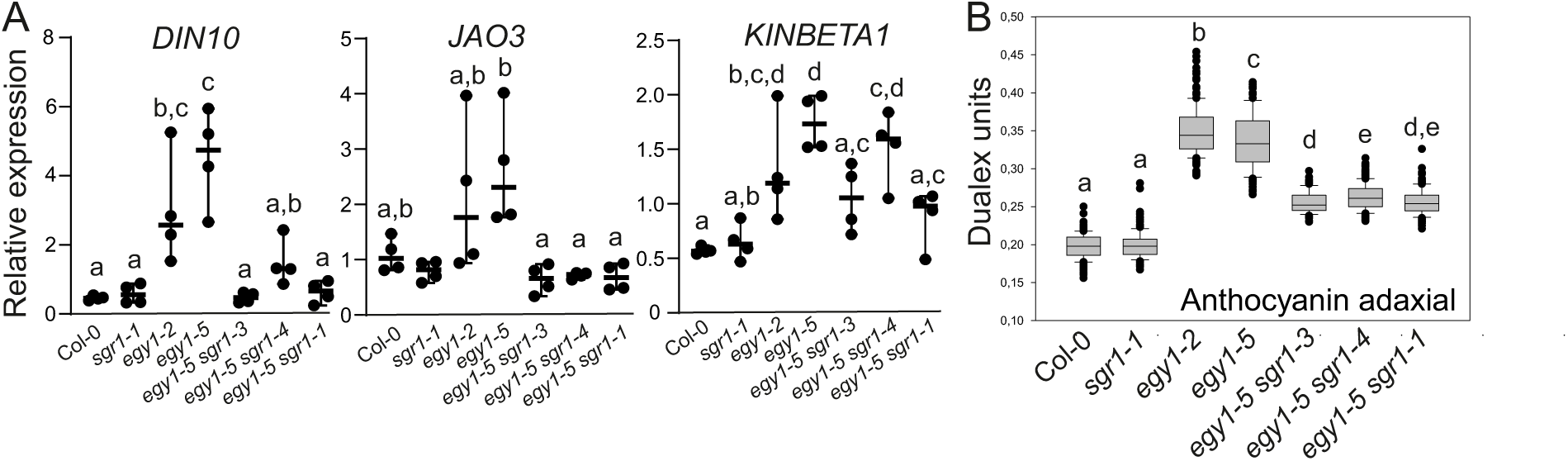
Retrograde signalling and anthocyanin accumulation in four-week-old plants. (A) Real-time quantitative PCR (qPCR) was used to assess the relative expression of *DIN10*, *JAO3* and *KINBETA1* normalized against three reference genes (*PP2AA3*, *TIP41* and *YLS8*). See Supplementary Fig. S8 for additional genes. Data represents four biological replicates. Genotypes with different letters are statistically different (1-way ANOVA and Tukey’s multiple comparisons test). (B) Anthocyanin content estimated with Dualex from ten biological repeats, N = 150 leaves. The box plot indicates the median, upper and lower quartile, and dots as outliers. Genotypes with different letters are statistically different (1-way ANOVA and Tukey’s multiple comparisons test).

Two marker genes involved in chloroplast retrograde signalling *LHCB2.4* (*LIGHT-HARVESTING CHLOROPHYLL B-BINDING 2.4*) and *VDEP* (*VIOLAXANTHIN DE-EPOXIDASE 1*) showed only minor changes at transcript levels (Supplementary Fig. S8). Retrograde signalling from the chloroplast also regulates biosynthesis of flavonoids and anthocyanin that protect the plant during light stress (Richter *et al*., 2023). We performed non-invasive measurements of anthocyanin content in leaves using Dualex. The anthocyanin content was significantly higher in *egy1-2* and *egy1-5* (Figure 5B), which suggests that there is an increased need to screen visible light to protect the chloroplast. The anthocyanin content was partially reduced to wildtype levels in the suppressor mutants, which is indicative of feedback mechanisms between chlorophyll breakdown and anthocyanin accumulation that are likely mediated by retrograde signalling.

### Chlorosis of *egy1* is downstream of the known senescence pathways

The breakdown of chlorophyll during plant senescence is a highly controlled process involving several identified regulators, including the plant hormones: salicylic acid (SA), jasmonic acid (JA), abscisic acid (ABA) and ethylene (Woo *et al*., 2019). In order to investigate whether the development of chlorosis in the *egy1* mutant was driven solely by events occurring within the chloroplast (i.e. chlorophyll breakdown), or whether signalling from outside the chloroplast was also contributing, we crossed the *egy1-5* mutant with the mutants deficient in regulators of hormone signalling or biosynthesis. These included *abi2* (*aba insensitive2*, impaired in ABA signalling (Leung *et al*., 1997)), *aos* (*allene oxide synthase*, a JA biosynthesis mutant (Park *et al*., 2002)), *coi1* (*coronatine insensitive1*, impaired in JA signalling (Ellis and Turner, 2002)), *eds1* (*enhanced disease susceptibility1*, defective in SA accumulation and pathogen signalling (Falk *et al*., 1999)), *ein2* (*ethylene insensitive2*, impaired in ethylene signalling (Alonso *et al*., 1999)), *rcd1* (*radical induced cell death1*, a pleiotropic mutant with defects in senescence regulation (Vainonen *et al*., 2012)), and *sid2* (*salicylic acid induction deficient2*, a SA biosynthesis mutant (Wildermuth *et al*., 2001)). In all cases the corresponding double mutants displayed the same phenotype as the single mutant *egy1-5* (Figure 6). Therefore, the chlorosis observed in the leaves of the *egy1-5* mutant is likely to be a downstream event of the known hormone-dependent senescence pathways.

**Figure 6.**
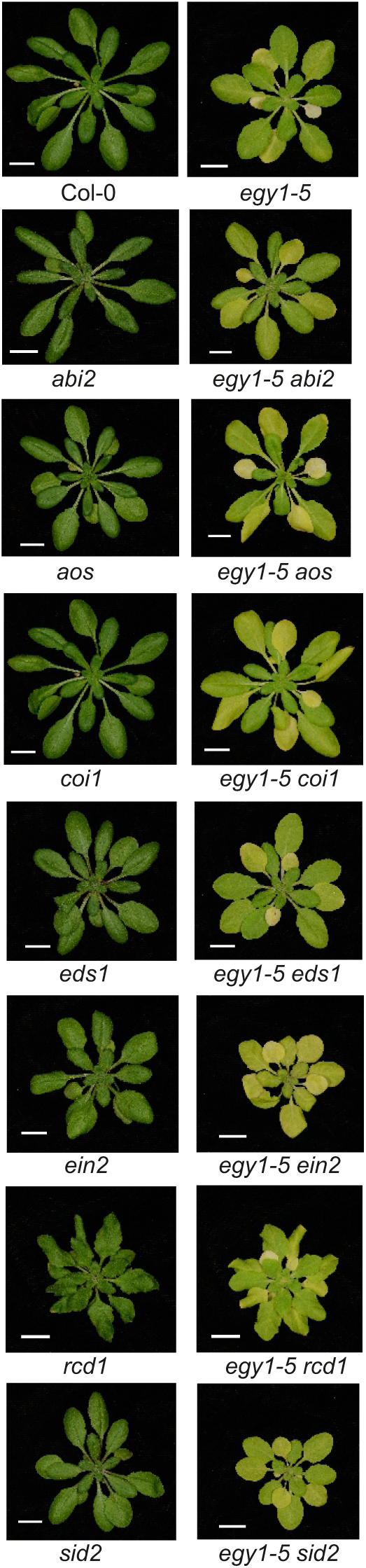
Double mutant analysis of *egy1-5* crossed to mutants defective in senescence regulation, hormone biosynthesis or signalling. Four-week-old plants of Col-0, *egy1-5*, *abi2-1*, *egy1-5 abi2-1*, *aos*, *egy1-5 aos*, *coi1-16*, *egy1-5 coi1-16*, *eds1-2*, *egy1-5 eds1-*2, *ein2-1*, *egy1-5 ein2-1*, *rcd1*, *egy1-5 rcd1*, *sid2-1*, *egy1-5 sid2-1*. Scale bar = 1 cm.

### Guard cell chloroplasts are absent from *egy1* and restored in *egy1 sgr1*

The *egy1-4* allele was recently used to demonstrate that EGY1 is necessary for guard cell chloroplast development (Sanjaya *et al*., 2021b). The guard cells of *egy1-4* did not have chloroplasts (Sanjaya *et al*., 2021b). This raised a question of whether EGY1 itself was required for guard cell chloroplast development. To address this, we used the suppressor mutants. If EGY1 were strictly required for guard cell chloroplast development, it would be expected that both *egy1* and *egy1 sgr1* lack guard cell chloroplasts. Conversely, if the absence of guard cell chloroplasts was the result of defective photosynthetic complexes, this should be restored in suppressor mutants. We monitored chlorophyll autofluorescence from leaves of 4–5-week-old plants using confocal microscopy (Figure 7). In line with a previous report (Sanjaya *et al*., 2021b), guard cells in *egy1-2* and *egy1-5* did not have chloroplasts; however, they were restored in *egy1-5 sgr1-3* and *egy1-5 sgr1-4* (Figure 7). This suggested that EGY1 is not a direct regulator of guard cell chloroplast development; rather, the failure of guard cell chloroplast development in *egy1* is due to deficient assembly of chlorophyll-binding complexes.

**Figure 7.**
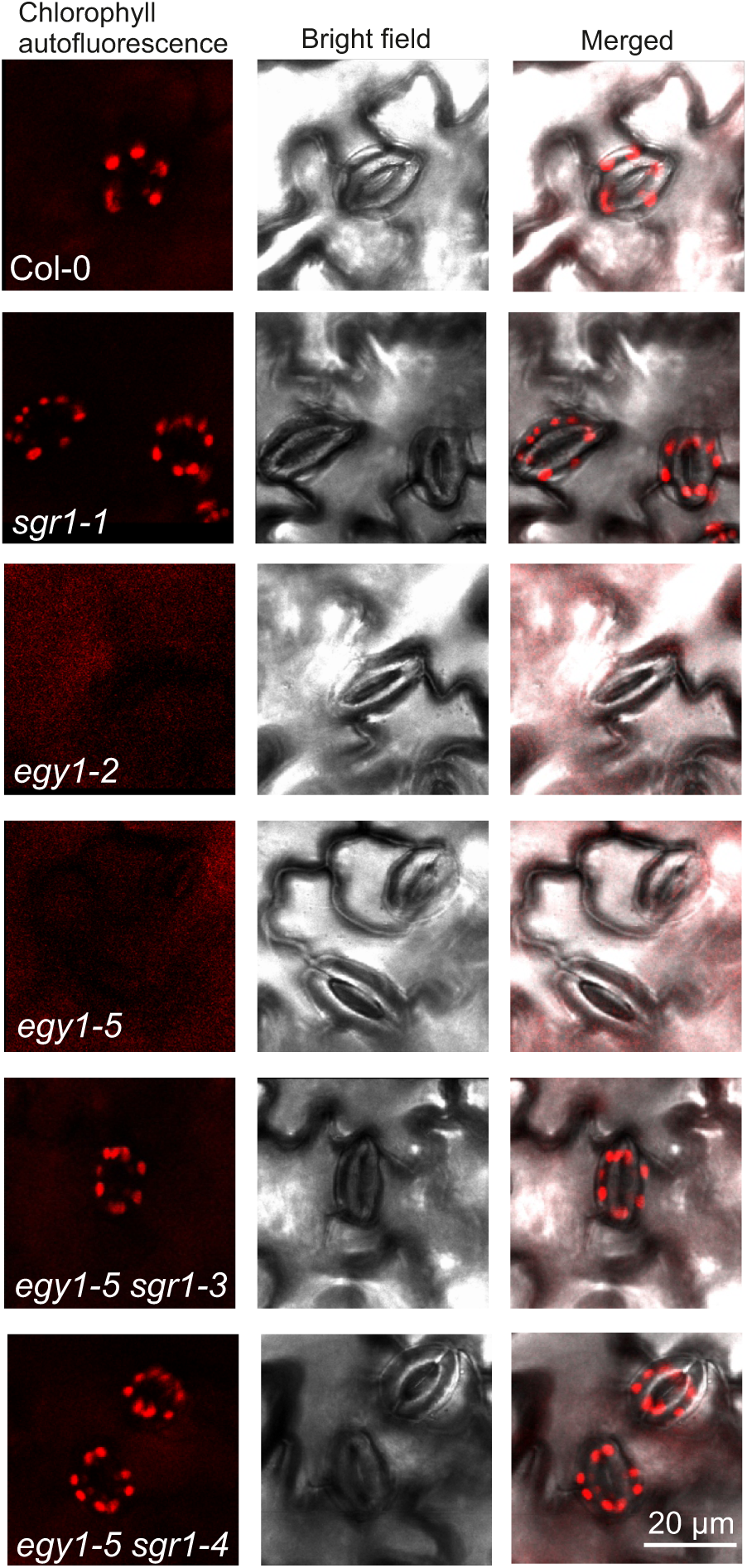
Guard cell chloroplasts are absent in *egy1* and restored in *egy1 sgr1*. Confocal microscopy images of fully developed four-week-old Arabidopsis leaves (abaxial side). Chlorophyll autofluorescence in guard cells is shown in red, bright field images and merged. Scale bar represents 20 μm.

## Discussion

This study describes the functional interaction between *EGY1* and *SGR1*. EGY1 is a thylakoid-associated protease and SGR1 is a chlorophyll *a*: Mg^2+^ dechelatase, which catalyses the early step of chlorophyll degradation. It has been shown that in a complex with BCM1, EGY1 can directly proteolyze SGR1, thereby controlling the biosynthesis and breakdown of chlorophyll (Fu *et al*., 2025). Here, we independently discovered the functional interaction between EGY1 and SGR1 using an unbiased forward genetic screen, and explored its implications for photosynthetic functions, cell signalling and chloroplast development.

### The impact of EGY1-SGR1 module on LHC proteins

At the chloroplast level, our results revealed novel interconnections between EGY1 and SGR1 in the biogenesis of photosynthetic complexes. The translation and chloroplast import of chlorophyll-binding proteins are tightly coordinated between the chloroplast and nuclear genomes, and are synchronized with chlorophyll metabolism (Wang and Grimm, 2021). The biosynthesis of chlorophyll occurs in strict stoichiometry with the availability of the subunits of the major chlorophyll-binding complex, LHCII, which are encoded by the nuclear genome. Loss of EGY1 function resulted in the depletion of all LHCII and LHCI subunits (Figure 3E, Table 1, (Fu *et al*., 2025)), though *LHCB2.4* transcript levels were not suppressed in *egy1* (Supplementary Fig. S8). Therefore, the depletion of LHCII and LHCI is unlikely due to repression at the transcriptional level, but rather to SGR1-dependent degradation of chlorophylls in LHC proteins (Tanaka and Ito, 2024). Indeed, in the *egy1 sgr1* suppressor mutants, abundance of LHCI and LHCII trimers as well as part of the PSII-LHCII complexes was restored to wildtype levels (Figure 3E, Supplementary Fig. S5).

Interestingly, while LHCII and LHCI bind most of the chlorophyll *b*, SGR1 only removes Mg from chlorophyll *a* (Sato *et al*., 2018). Accordingly, genetic perturbations of *EGY1* and *SGR1* affect cellular chlorophyll *a/b* ratios (Figure 2), consistent with the significant depletion of LHC in *egy1* compared to other chlorophyll-binding proteins (Figure 3E, Supplementary Fig. S5 and (Adamiec *et al*., 2021)). Accumulation of SGR1 stimulates the expression of the chlorophyll *b* reductase NON-YELLOW COLOURING 1 (NYC1) (Sato *et al*., 2018), and promotes the propagation of senescence (Tanaka and Ito, 2024). Therefore, the depletion of the chlorophyll *b*-binding proteins LHCII and LHCI is likely caused by the constitutive accumulation of SGR1 in *egy1* (Fu *et al*., 2025), which stimulates the conversion of chlorophyll *b* to *a* by NYC1. Consistent with this, chlorophyll *a/b* ratios in the *egy1 sgr1* mutants were similar to those in the wildtype, suggesting that in *sgr1* the changes in chlorophyll *a* abundance directly influence the abundance of chlorophyll *b* through the chlorophyll *a*-to-*b* conversion systems (Wang and Grimm, 2021). Our results support the idea that SGR1 is also expressed in pre-senescent leaves, but this expression is undetectable due to the control exerted by the BCM1-EGY1 module (Fu *et al*., 2025; Yamatani *et al*., 2022). Furthermore, chlorophyll degradation is also involved in the degradation of LHCII under prolonged exposure to high light (Sato *et al*., 2015), suggesting that the control of SGR1 by the BCM1-EGY1 module is not only limited to senescence, but extends to acclimation to light intensity as well. This may explain the higher amounts of chlorophyll *a* and *b* in *sgr1* (Figure 2C). The lack of EGY1 allows increased SGR1 accumulation from early stages, leading to progressive self-activation of senescence during growth (Tanaka and Ito, 2024). This explains the deterioration of the senescence phenotype of *egy1* with aging (Figure 1-3 and Supplementary Fig. S5). SGR1-dependent chlorophyll *a* degradation induced by senescence affects PSI (Chen *et al*., 2021b), which is consistent with the depletion of PSI in *egy1* (Fu *et al*., 2025), a phenomenon that is reverted in *egy1 sgr1* (Figure 3, Supplementary Figs. S5 and S6). The depletion in PSI in *egy1* may be linked to high sensitivity of the mutant to a photosynthetic inhibitor MV (Figure 4). However, MV toxicity depends on many factors, including cellular redox balance, the functions of chloroplast thiol redox systems and mitochondrial respiration (Shapiguzov *et al*., 2019). The mechanisms underlying the sensitivity of *egy1* to MV require further investigation.

LHCII plays a key role in forming stacked thylakoid membranes (grana), inducing strict lateral heterogeneity in the distribution of PSII and PSI (Trotta *et al*., 2025). Previous analyses of thylakoid ultrastructure in *egy1* mutants revealed a significantly reduced level of thylakoid stacking (Chen *et al*., 2005; Sanjaya *et al*., 2021a; Sanjaya *et al*., 2021b). Therefore, it is plausible that the depletion of LHCII results in the unstacked thylakoids observed in *egy1* mutants, leading to loss of PSII and PSI lateral heterogeneity and, as a result, increased transfer of excitation energy from PSII to PSI (Trotta *et al*., 2025). This could explain the higher Fv/Fm ratio observed in *egy1* mutants when treated with lincomycin (Supplementary Fig. S7), indicating that PSI quenches PSII excitation in *egy1* mutants, thereby preventing PSII damage at low light intensities. Restoring LHCII abundance in *egy1 sgr1* enables stacked membranes and lateral heterogeneity to form, which may restrict PSII-to-PSI energy transfer and lead to a wildtype response of Fv/Fm under lincomycin treatment (Supplementary Fig. S7).

The inability of the *egy1 sgr1* and *egy1 sgr1 sgr2* mutants to suppress all *egy1* phenotypes suggests the existence of additional chlorophyll degradation mechanisms. Several other components have been proposed to play a role in this process, including STAY-GREEN LIKE (SGRL) (Chen *et al*., 2021b). Interestingly, the degradation of LHCII and PSI was suppressed in the *sgr1/sgr2/sgrl* triple mutant, while PSII degradation remained largely unaffected (Chen *et al*., 2021b). The role of EGY1 in various chlorophyll-degrading systems and its impact on the biogenesis of different thylakoid complexes remains to be elucidated.

### The potential role of EGY1 in photosystem assembly

Complete suppression of the LHC depletion but not of photosystem assembly in the *egy1 sgr1* double mutant indicates that the EGY1-SGR1 module degrades SGR1 in a manner that is limited to controlling the LHCs. This suggests an overlooked role for EGY1. Indeed, the double mutant *egy1 sgr1* still accumulates PSII and PSI assembly intermediates (Figure 3E, Supplementary Figs. S5 and S6) and exhibits altered chlorophyll fluorescence dynamics (Figure 3). Therefore, EGY1 may affect photosystem assembly independently of SGR1 activity. Several pieces of evidence point in this direction. During de-etiolation and thylakoid biogenesis, the absence of EGY1 affects the expression of LHC proteins but not that of D1 (Qi *et al*., 2020). However, mature *egy1* plants exhibit over-accumulation of D1 and depletion of CP47 and D2 (Adamiec *et al*., 2021), as well as depletion of several PSI subunits (Supplementary Fig. S6; (Fu *et al*., 2025)), suggesting a role for EGY1 in coordinating the expression and assembly of photosystem subunits.

EGY1 belongs to the S2P family of conserved proteases, four of which (EGY1, EGY2, EGY3 and S2P2) are thylakoid-localized in Arabidopsis (Adamiec *et al*., 2017). Mutations in these proteases affect the steady-state levels of LHC proteins (EGY1 and S2P2), PSII subunits (EGY2 and SP2P), PSI subunits (EGY1), and the degradation rates of PSII (EGY1 and EGY2) and PSI and LHC subunits (EGY3) under HL treatment (Figure 3, Supplementary Figs. S5 and S6; (Adamiec *et al*., 2022; Adamiec *et al*., 2021; Adamiec *et al*., 2020; Adamiec *et al*., 2018; Chen *et al*., 2021a; Chen *et al*., 2005; Fu *et al*., 2025)). However, a lack of EGY1 is the only case in which PSII assembly complexes accumulate (Figure 3, Supplementary Figs. S5 and S6; (Adamiec *et al*., 2021; Qi *et al*., 2020)). Furthermore, EGY2 interacts with components of the chloroplast transcriptional machinery, and the mRNA of specific chloroplast-encoded PSII subunits is depleted in *egy2* (Adamiec *et al*., 2018; Lucinski *et al*., 2024). Interestingly, the S2P family of proteases is involved in the degradation of membrane-associated transcription factors (Rudner *et al*., 1999). Both EGY1 and EGY2 have been suggested to degrade transcription factors associated with plastid RNA polymerase (Chen *et al*., 2012; Chen *et al*., 2021b). All S2P thylakoid proteases are localized in stroma-exposed membranes (Trotta *et al*., 2025), where the thylakoid-associated translational machinery resides. The latter is involved in replacing the damaged PSII D1 subunit under light conditions. Considering the above, and that EGY1 interacts with PSII subunits (Chen *et al*., 2021b), it is likely that the interaction between EGY1 and PSII (and possibly PSI) is directly linked to photosystem assembly.

In addition to SGR1, EGY1 together with BCM1, interact with GENOMES UNCOUPLED 4 (GUN4) and the CHLH/GUN5 components of the Mg chelatase complex, which are responsible for forming the tetrapyrrole intermediate Mg-protoporphyrin IX (Fu *et al*., 2025; Wang *et al*., 2025). However, these interactions do not appear to involve the proteolytic activity of EGY1; rather, EGY1 may function as a regulatory scaffold connecting tetrapyrrole biosynthesis with photosystem assembly. This possible second role of EGY1 is consistent with the observation that the *egy1-4* allele still accumulates PSII-LHCII supercomplexes (Qi *et al*., 2020). This is further reinforced by the fact that BCM1 and CHLH accumulate in both *egy1* and *ftsh2* mutants, but not in *clpc* mutants, which lack the ATP-binding subunit of the ATP-dependent Clp protease. Furthermore, EGY1 accumulates in *ftsh2* and *clpc* (Fu *et al*., 2025), and FTSH levels are higher in *egy1* background (Adamiec *et al*., 2021), revealing an interdependency between EGY1 and other proteases involved in photosystem assembly.

Taken together, the assembly of wildtype photosynthetic supercomplexes was not fully restored in *egy1 sgr1*. Native gels revealed that the intermediate assembly complexes of PSI (PSI*) and PSII (PSII monomer) were still more abundant in *egy1 sgr1* lines than in Col-0. This finding was corroborated by chlorophyll fluorometry assays, which revealed that the *egy1 sgr1* suppressors exhibited only partial complementation of wildtype functions (Figure 3). Overall, the results suggest that EGY1 is involved in the biogenesis of both LHCII and photosystem core components, whereas the SGR1 module only contributes to the biogenesis of LHCII.

### The impact of the EGY1-SGR1 module on signalling and development

The plant hormones ethylene, ABA, JA, and SA are essential regulators of both senescence and chlorophyll degradation (Guo *et al*., 2021). However, introducing mutations that disrupt hormone signalling or biosynthesis did not affect the development of leaf chlorosis in the *egy1* mutant (Figure 6). This suggests that the chlorosis observed in *egy1* occurs downstream of hormone signalling and is driven by processes from inside the chloroplast, such as chlorophyll homeostasis and the assembly of photosynthetic complexes, as discussed above (Figure 8).

**Figure 8.**
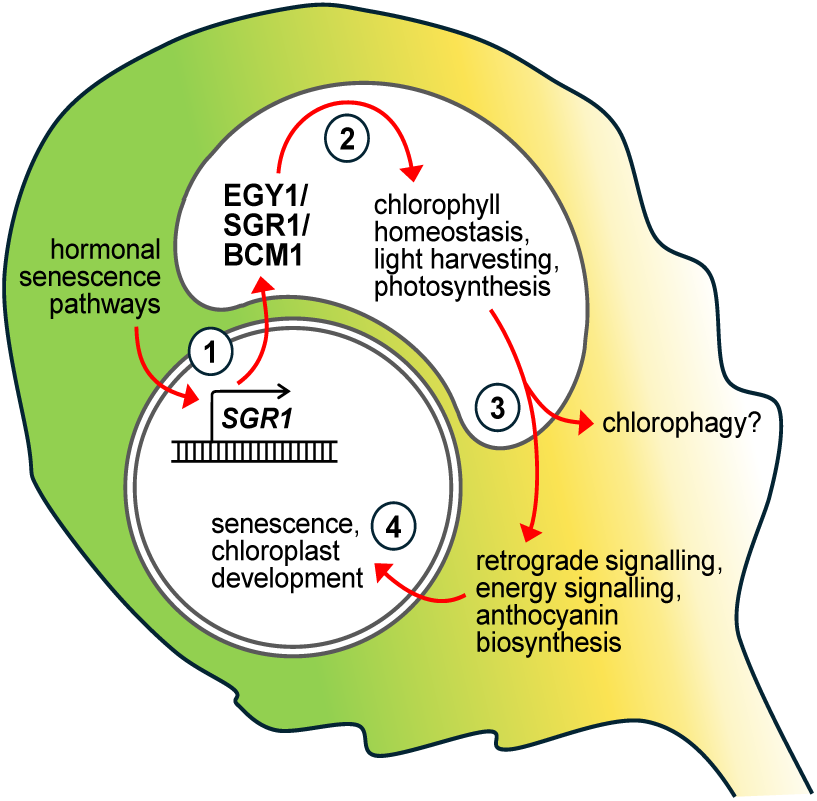
The EGY1-SGR1 regulatory module controls several aspects of chloroplast biology. **1**. Upstream regulators of senescence include hormonal signalling pathways that directly target *SGR1* and other chlorophyll catabolism genes. **2**. Inside the chloroplast, the EGY1/SGR1/BCM1 regulatory complex balances chlorophyll biosynthesis and catabolism. When EGY1 is absent in the *egy1* mutant, this leads to adverse chlorophyll homeostasis which in turn results in incorrect assembly of photosynthetic protein complexes and to abnormal photosynthetic electron transfer. These defects are partially or fully restored when chlorophyll levels are increased through mutations in *sgr1*. **3**. In *egy1* guard cells, the chloroplasts are absent. Either they fail to develop, or defective chloroplasts are removed through processes like chlorophagy. Importantly, as guard cell chloroplasts are restored in *egy1 sgr1*, this shows that it is not EGY1 that directly regulates chloroplast development, rather it is the defects in photosynthetic complexes that likely initiate the removal of chloroplasts. **4**. Various signals leave the chloroplast to co-ordinate gene expression with the nucleus. This includes regulation of anthocyanin biosynthesis, which is also under control of the EGY1-SGR1 module.

Few of the tested marker genes related to retrograde signalling showed major transcriptional changes (Figure 5). *DIN10* was highly induced in *egy1* and restored to wildtype levels in *egy1 sgr1*. This gene encodes RAFFINOSE SYNTHASE 6. The trisaccharide raffinose is proposed to protect against various abiotic and biotic stresses (Yan *et al*., 2022). *DIN10* was originally identified in a screen for transcripts induced by dark treatment; however, *DIN10* transcripts were also suppressed by sucrose treatment (Fujiki *et al*., 2001). This suggests that expression of *DIN10* can act as a proxy for the energy (sugar) status of the chloroplast. This could explain why *DIN10* transcript levels are elevated in *egy1* and suppressed in *egy1 sgr1*. In *egy1*, the constitutive accumulation of SGR1 (Fu *et al*., 2025), or the impaired chloroplast function, may activate retrograde signalling leading to increased *DIN10* expression. Indeed, when SGR1 function is impaired in suppressor mutants *egy1-5 sgr1*, the *DIN10* transcript levels return to wildtype levels (Figure 5).

The EGY1-SGR1 regulatory module plays a key role in regulating multiple processes related to the chloroplast (Figure 8). Upstream of the chloroplast, senescence signalling mediated by hormones regulates the transcription of senescence-associated genes, including *SGR1*. Within the chloroplast, the EGY1/SGR1/BCM1 protein complex coordinates chlorophyll metabolism (Fu *et al*., 2025). Reduced chlorophyll levels (as observed in *egy1*) lead to defects in the assembly of photosynthetic supercomplexes and defects in photosynthetic electron transfer (Figure 3). These defects result in the activation of retrograde signalling and subsequent transcriptional reprogramming, causing, among other consequences, the activation of anthocyanin biosynthesis. It has been proposed that EGY1 is a regulator of chloroplast development (Chen *et al*., 2005; Sanjaya *et al*., 2021a; Sanjaya *et al*., 2021b). In *egy1* guard cells, no chlorophyll fluorescence was observed, indicating a complete absence of functional chloroplasts (Figure 7, (Sanjaya *et al*., 2021b)). However, restoration of guard cell chloroplasts in *egy1 sgr1* revealed that EGY1 does not directly regulate guard cell chloroplast development. Instead, we propose that the improper assembly of chloroplast protein complexes in *egy1* (Figure 3E) damages the chloroplasts, resulting in their removal through processes such as chlorophagy (Figure 8, (Nakamura and Izumi, 2018)). This sequence of events is corroborated by the increased sensitivity of *egy1* to chemicals that generate reactive oxygen species (Figure 4). Improved chlorophyll homeostasis in *egy1 sgr1* restores photosynthetic electron transfer, stabilizes protein complexes, normalizes retrograde signalling, and ultimately re-establishes functional guard cell chloroplasts.

## Material and methods

### Plant material and growth conditions

The plants were grown in 1:1 peat-vermiculite under 250 µmol m^2^ s^−1^ white light irradiance, 12 h : 12 h light-dark cycle, 23°C/19°C (day/night) temperature and 70%/90% relative humidity. Unless stated otherwise, 4-week-old plants used for experiments. The spontaneous *egy1-5* mutant was identified among Col-0 plants during routine experimental procedures. Other mutants were obtained from stock centre (https://arabidopsis.info/) *aos, coi1-16, ein2-1*, *rcd1-4, sid2-1* or were gifts, *egy1-2* (Prof. Ning Li (Chen *et al*., 2005)), *abi2-1* (Prof. Julian Schroeder), *eds1-2* (Prof. Jane Parker), *sgr1-1*, *sgr2* and *sgr1 sgr2* (Prof. Nam-Chon Paek (Sakuraba *et al*., 2014)).

### Identification of egy1-5

Two approaches were used to identify the causative mutation. The new mutant was crossed to Ler to generate a F2 population for traditional mapping using simple sequence length polymorphism markers (Supplementary Table S2). This showed strong linkage to 11-12 Mbp on chromosome 5. Subsequently whole genome sequencing identified a four bp deletion in the gene *EGY1* (AT5G35220), Supplementary Figure 1. The four bp deletion was further confirmed by Sanger sequencing of the *EGY1* cDNA from *egy1-5*. A derived cleaved amplified polymorphic sequence (dCAPS) marker was developed to detect presence of the *egy1-5* deletion (Supplementary Table S2). For allelism test, *egy1-5* was crossed to the T-DNA allele *egy1-2* (SALK_134931), and F1 plants were evaluated for the senescence phenotype and genotyped with *egy1-5* dCAPS marker and with *egy1-2* T-DNA genotyping primers (Supplementary Table S2).

### Suppressor mutant screen and identification of new stay green1 mutants

Seeds from *egy1-5* were mutagenized with EMS according to (Weigel and Glazebrook, 2006). Approximately 10 000 M2 seeds were planted at high density and screened for plants with restored green colour in leaves of 4-5 week old plants. Seeds from the best suppressors were re-screened in M3 and the two best suppressors (S1 and S2) used for further studies. S2 was backcrossed to *egy1-5* to generate a mapping population. Using 600 F2 plants, leaves from 140 plants with the suppressor phenotype were pooled for genomic DNA isolation and mapping of the causative mutation with ShoreMap 3.0, (Sun and Schneeberger, 2015), Supplementary Fig. S1. This identified a G to A mutation at position 452 in the coding sequence for *STAY GREEN1* (*SGR1*) (AT4G22920), corresponding to R151Q in the protein. Sanger sequencing of STAY GREEN1 in both suppressors confirmed the G to A mutation in suppressor 2, and in suppressor 1 identified a G to A mutation at position 472 of the coding sequence of SGR1, corresponding to E158K in the protein. The new mutant alleles for *sgr1* were renamed *sgr1-3* (suppressor 1) and *sgr1-4* (suppressor 2) and PCR based markers were developed for genotyping (Supplementary Table S2).

To confirm that mutations in *sgr1* suppress *egy1* phenotypes, an independent mutant allele for *sgr1* (*sgr1-1*, originally known as *nye1*, (Ren *et al*., 2007)) were crossed with *egy1-5* to generate *egy1-5 sgr1-1*. The *sgr1-1* mutation was genotyped with dCAPS marker (Supplementary Table S2). The *egy1-5* allele was also crossed to *sgr1-1 sgr2* and to *sgr2* (Sakuraba *et al*., 2014), to generate the double mutant *egy1-5 sgr2* and the triple mutant *egy1-5 sgr1-1 sgr2*.

Suppressors 1 and 2 were complemented by transformation with the wildtype STAY GREEN1 gene including 1.1 kb promoter region. SGR1 was amplified from Col-0 DNA with primers containing attB sites for Gateway cloning (Supplementary Table S2). PCR products were first introduced to pDONR-Zeo and after sequencing transferred to pMDC100, introduced to Agrobacterium GV3101 and transformed to suppressors 1 and 2 via floral dip (Clough and Bent, 1998).

### Double mutants

Crosses were made with *egy1-5* as pollen acceptor, and *abi2-1*, *eds1-2*, *ein2-1*, *rcd1-4* and *sid2-1* as pollen donors. For hormone mutants that are partially (*coi1-16*) or fully (*aos*) male sterile, these were used as pollen acceptors and *egy1-5* as the pollen donor. In addition, *egy1-5* and *var2* was crossed with *sgr1-1 sgr2*. Homozygous F2 plants were identified with PCR based markers (Supplementary Table S2).

### RNA isolation and real time quantitative PCR

RNA was isolated using GeneJET Plant RNA Purification Kit (ThermoFisher Scientific), from four week old plants. RNA (1 µg) was treated with DNase I (ThermoFisher Scientific) and used for cDNA synthesis with Maxima Reverse Transcriptase (ThermoFisher Scientific). After cDNA synthesis, final volume was diluted to 100 µl, and 1 µl used for qPCR with 5X EvaGreen qPCR mix (Solis Biodyne). Reactions were run on CFX384 Touch Real-Time PCR Detection System (Bio-Rad). Primers used for qRT–PCR, and their amplification efficiency are listed in (Supplementary Table S2). Analysis of qPCR results was performed using qBase3.4 (CellCarta), (Hellemans *et al*., 2007). Three reference genes (*PP2AA3*, *TIP41*, *YLS8*) were used for normalization.

### Measurement of Chlorophyll and other pigments

Two methods were used to measure chlorophyll content: Dualex and extraction chlorophyll with acetone for spectrophotometric measurement of chl *a* and chl *b*. With Dualex Scientific + (FORCE-A) the chlorophyll content and epidermal flavonols and anthocyanins are obtained per area basis. For acetone extraction of chl *a* and chl *b*, a single leaf of the same developmental stage (leaf number 5) from each genotype was weighted and put to 1.5 ml Eppendorf tubes with glass beads and frozen in liquid nitrogen. The leaf was broken into powder by shaking in a Silamat S6 (Ivoclar vivadent), followed by extraction in 80% acetone. Absorption was measured at 645 nm and 663 nm in a Nanodrop spectrophotometer.

### Physiological analyses of photosynthesis

OJIP kinetics was recorded using FluorPen FP 100 (Photon System Instruments, Drásov, Czech Republic). For measurements of chlorophyl fluorescence Kautsky kinetics, plants were acclimated to darkness for at least 30 min, then exposed to low-intensity (40 µmol photons m^−2^ s^−1^) blue light (450 nm) inside the IMAGING-PAM M-Series (Walz, Effeltrich, Germany). For photoinhibition assays, leaf discs were soaked overnight in H2O supplemented with 0.05% (v/v) Tween-20 with or without added MV, lincomycin. Then photoinhibition was assayed using IMAGING-PAM essentially as described in (Shapiguzov and Kangasjarvi, 2022). Non-photochemical quenching (NPQ) was measured using IMAGING-PAM. Plants were dark acclimated for at least 30 min, then minimal (Fo) and maximal (Fm) fluorescence were determined. Next, low-intensity actinic light (450 nm, approximately 40 µmol m^−2^ s^−1^) was turned on for 14 min, after which a saturating light pulse was triggered to measure light-adapted maximal fluorescence (Fm’). Then actinic light intensity was increased to approximately 230 µmol m^−2^ s^−1^ for 14 min, followed by the second Fm’ measurement. For each light intensity level, NPQ was calculated as: NPQ = (Fm−Fm’)/Fm’ (Horton and Ruban, 1992).

### Biochemical analyses of photosynthetic complexes

For blue-native and 2-dimentional gel analyses, thylakoids were isolated from leaf material as described in (Järvi *et al*., 2011), solubilized and separated by gel electrophoresis essentially as described in (Suorsa *et al*., 2015). The L-LHCII bands resolved by lpBN-PAGE from thylakoids isolated from Col-0, *egy1-5* and *egy1-5 sgr1-3* grown under long day conditions (16h light/8h darkness) (Table 1, Supplementary Table S1) were analysed by mass spectrometry-based proteomics as described in Gerotto et al. 2022.

### Chlorophyll autofluorescence

To detect chlorophyll autofluorescence, epidermal layer of fully developed 4-week old Arabidopsis leaves (plants grown as indicated above), were imaged with a Leica Stellaris 8 confocal microscope equipped with the Leica Application Suite X package (water immersion). For chlorophyll autofluorescence, excitation was performed using a 488 laser, emission 630–700 nm. Experiments performed in three biological repeats, confocal settings were kept identical in the presented samples.

### Statistics

Statistical tests were performed in GraphPad Prism version 10.4.2. Statistical parameters such as box plots are indicated in the figure legends. In the graphs, letters indicate statistical significance tests by analysis of variance (ANOVA). Sample numbers and biological replicates are indicated for all data points in figure captions.

## Data availability

Sequence data is available from the Sequence Read Archive under the accession number PRJNA1310757.

## Acknowledgements

This project was funded by the Research council of Finland (349540 and 363290 to M.B.; 346140 to A.S.), the Jane and Aatos Erkko Foundation (A.T. and E-M.A) and Estonian Research Council (T.V. grant nr: STP51). We thank Leena Grönholm for assistance during greenhouse experiments, Tuomas Puukko for assistance with mutant mapping and photography, and Maija Sierla for comments on the manuscript.

---

**Fig. S1.**
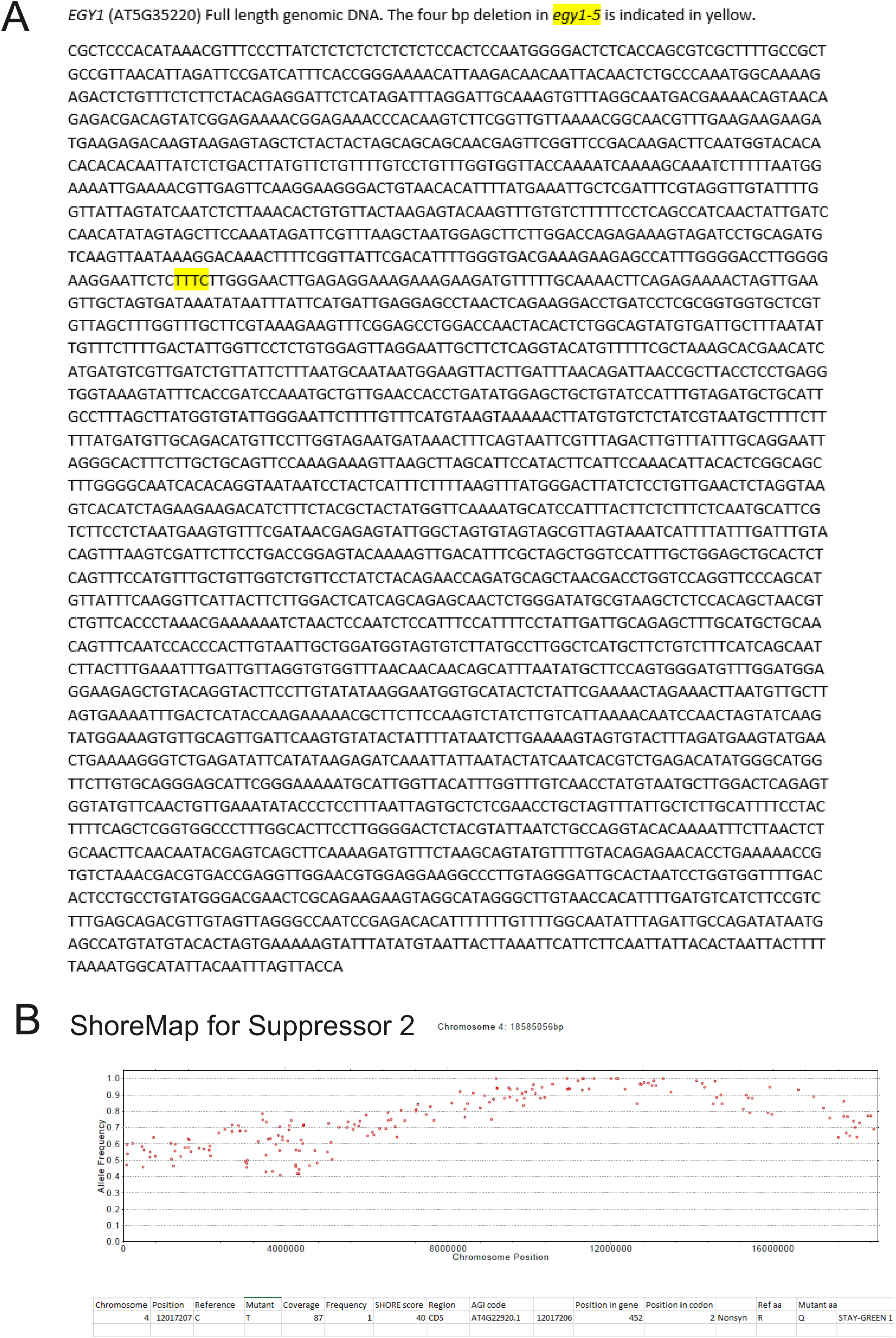
(A) Location of the four base pair deletion in *egy1-5* is indicated with yellow highlight. (B) ShoreMap results for suppressor 2 (*egy1-5 sgr1-4*), identified a point mutation in *SGR1* (*AT4G22920*) corresponding to a G to A mutation at position 452. The gene for SGR1 is located on the reverse strand on chromosome 4, hence the ShoreMap results shows this mutation as a C to T mutation.

**Fig. S2.**
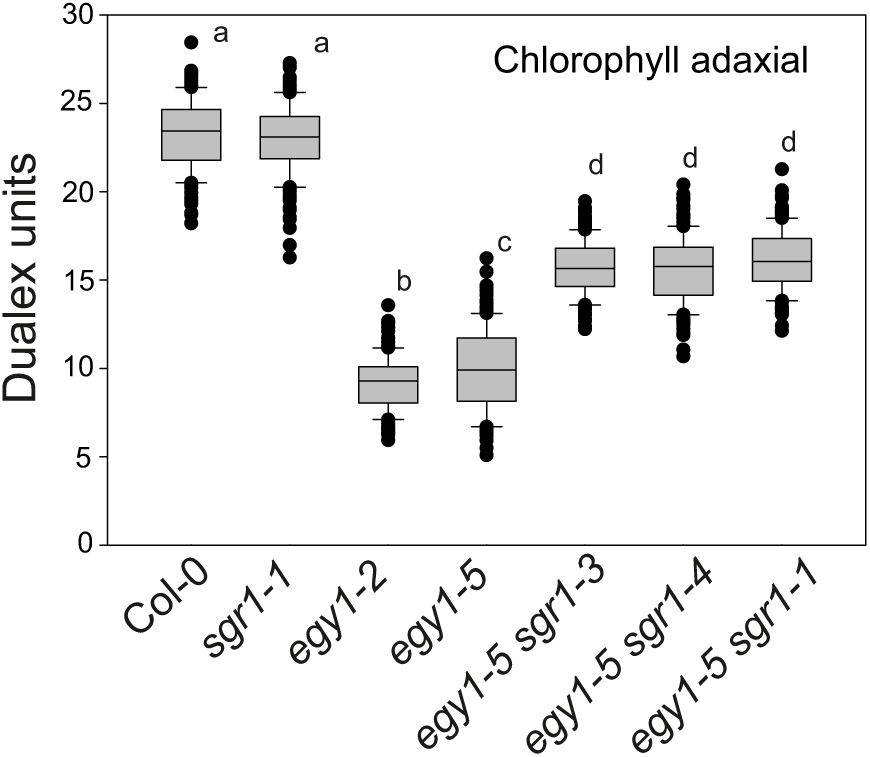
Chlorophyll content *in vivo* measured with Dualex in four-week-old plants, from ten biological repeats, N = 150 leaves. The box plot indicates the median, upper and lower quartile, and dots as outliers. Genotypes with different letters are statistically different (1-way ANOVA and Tukey’s multiple comparisons test).

**Fig. S3.**
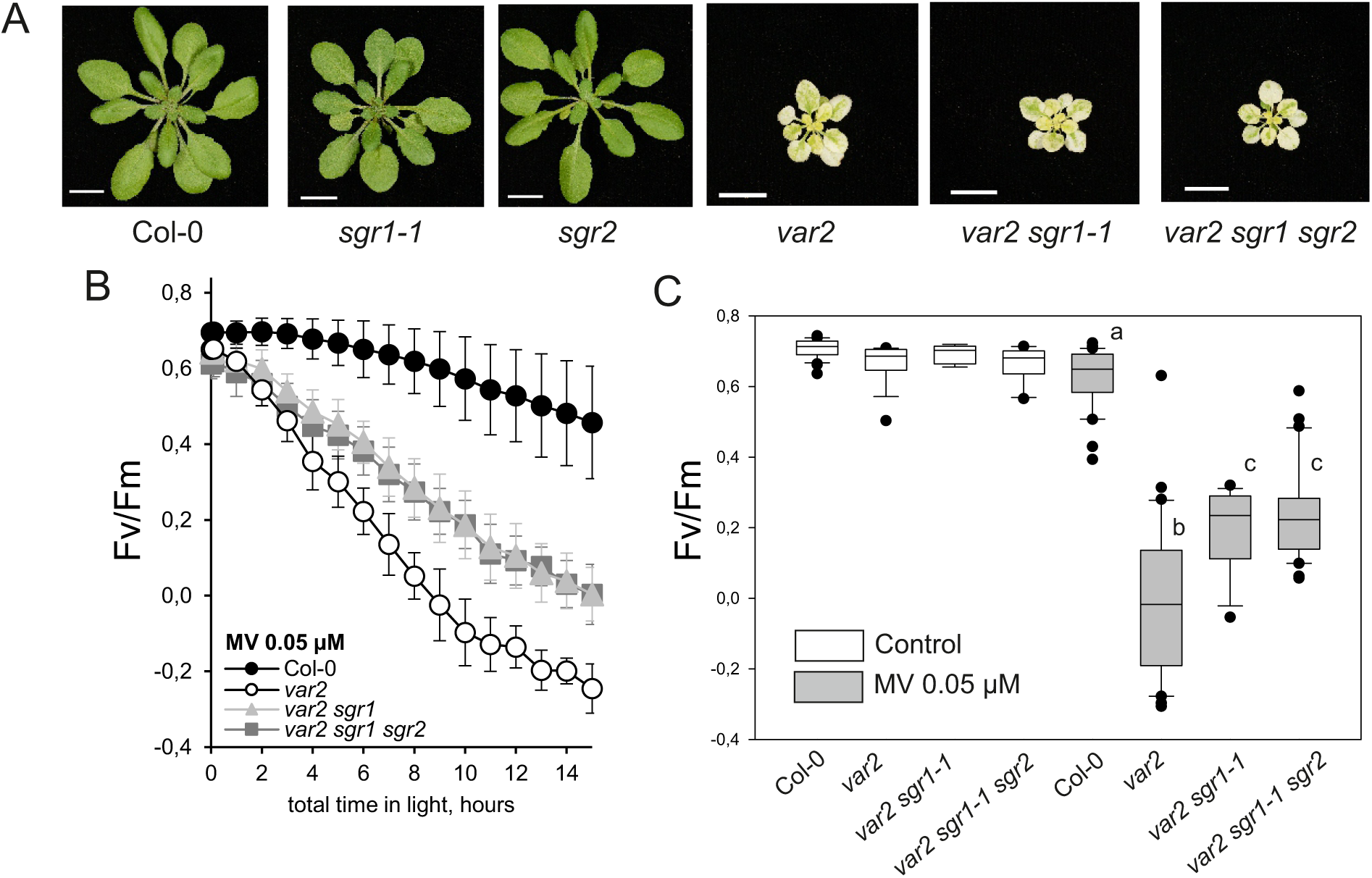
The *sgr1* mutant partially rescues MV sensitivity of *var2*, but not the visible variegated phenotype. (A) Four-week-old plants of Col-0, *sgr1*, *sgr2*, *var2*, *var2 sgr1* and *var2 sgr1 sgr2*. Scale bar = 1 cm. (B) Fv/Fm measured in leaf discs exposed to repeated cycles of blue light (448nm, 80 µmol m^−2^ s^−1^) and 0.05 µM MV. N = 5 leaf discs, the error bars indicate the SD. ***C***, Data from the 10 hr timepoint was quantified from three biological repeats, N=15-30 leaf discs. The box plot indicates the median, upper and lower quartile, and dots as outliers. Genotypes with different letters are statistically different (1-way ANOVA and Tukey’s multiple comparisons test).

**Fig. S4.**
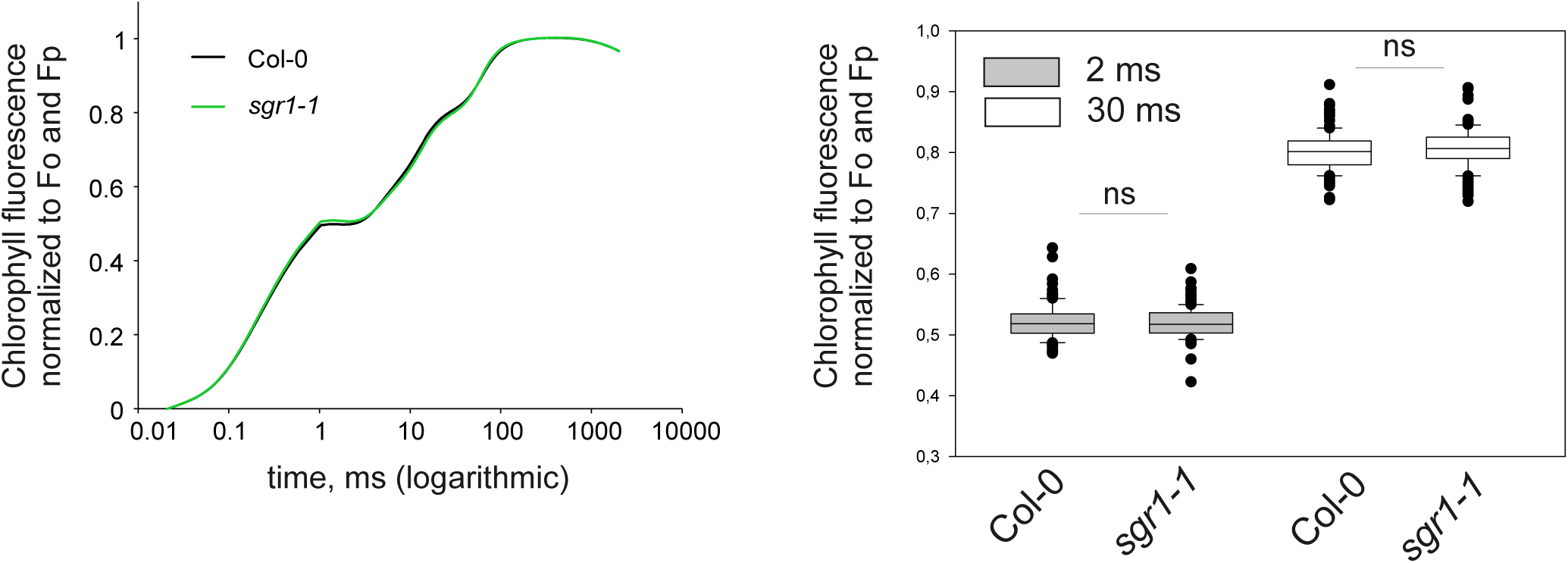
OJIP kinetics in four-week-old plants of Col-0 and *sgr1-1*. OJIP was double normalized to Fo and Fp, and data was extracted for the 2 and 30 ms time points from six biological repeats, N = 120 leaves. No statistically significant difference was detected between Col-0 and *sgr1-1* (Student t-test).

**Fig. S5.**
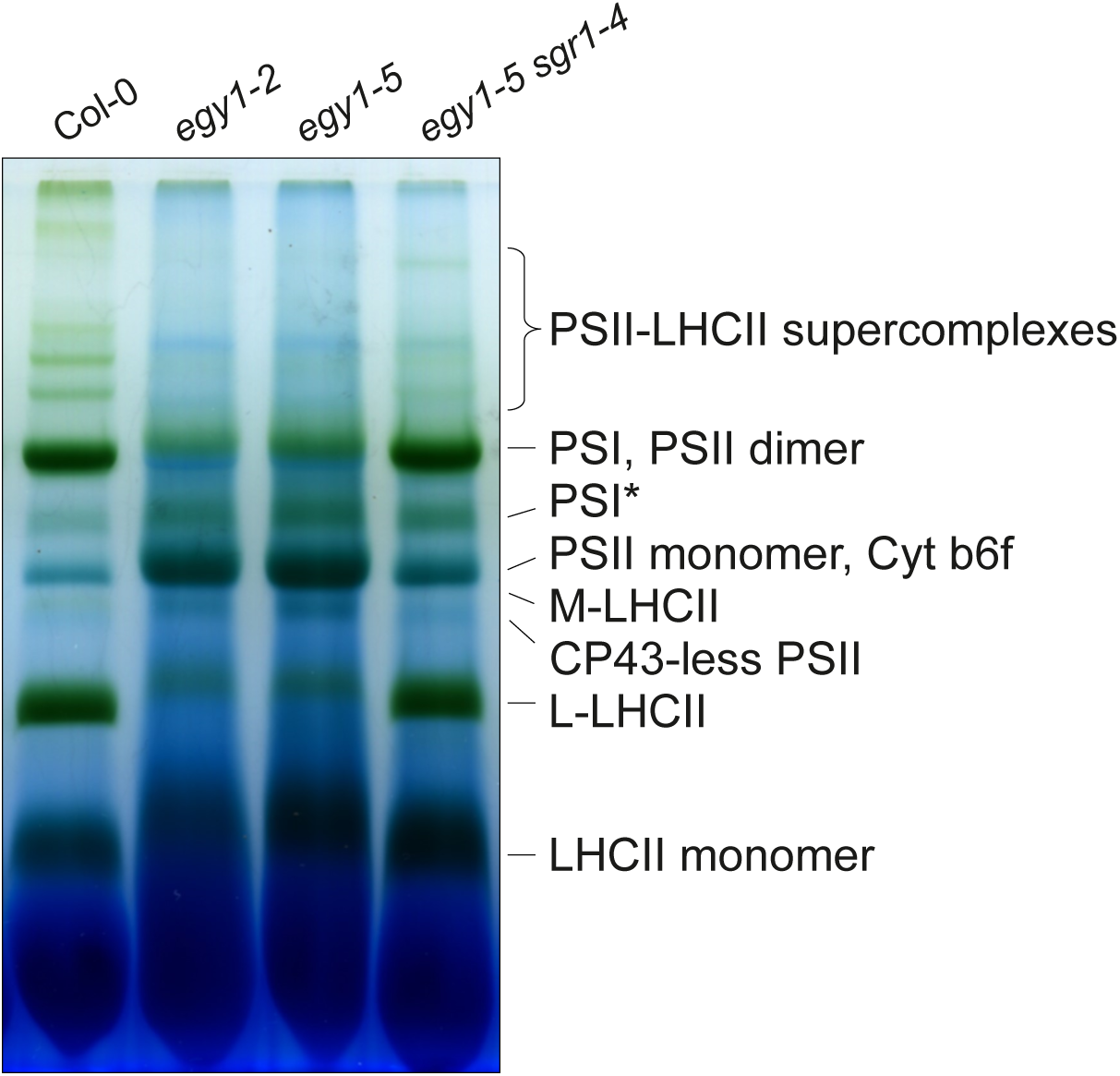
Separation by lpBN-PAGE of protein complexes from thylakoids isolated from 35 days old plants.

**Fig. S6.**
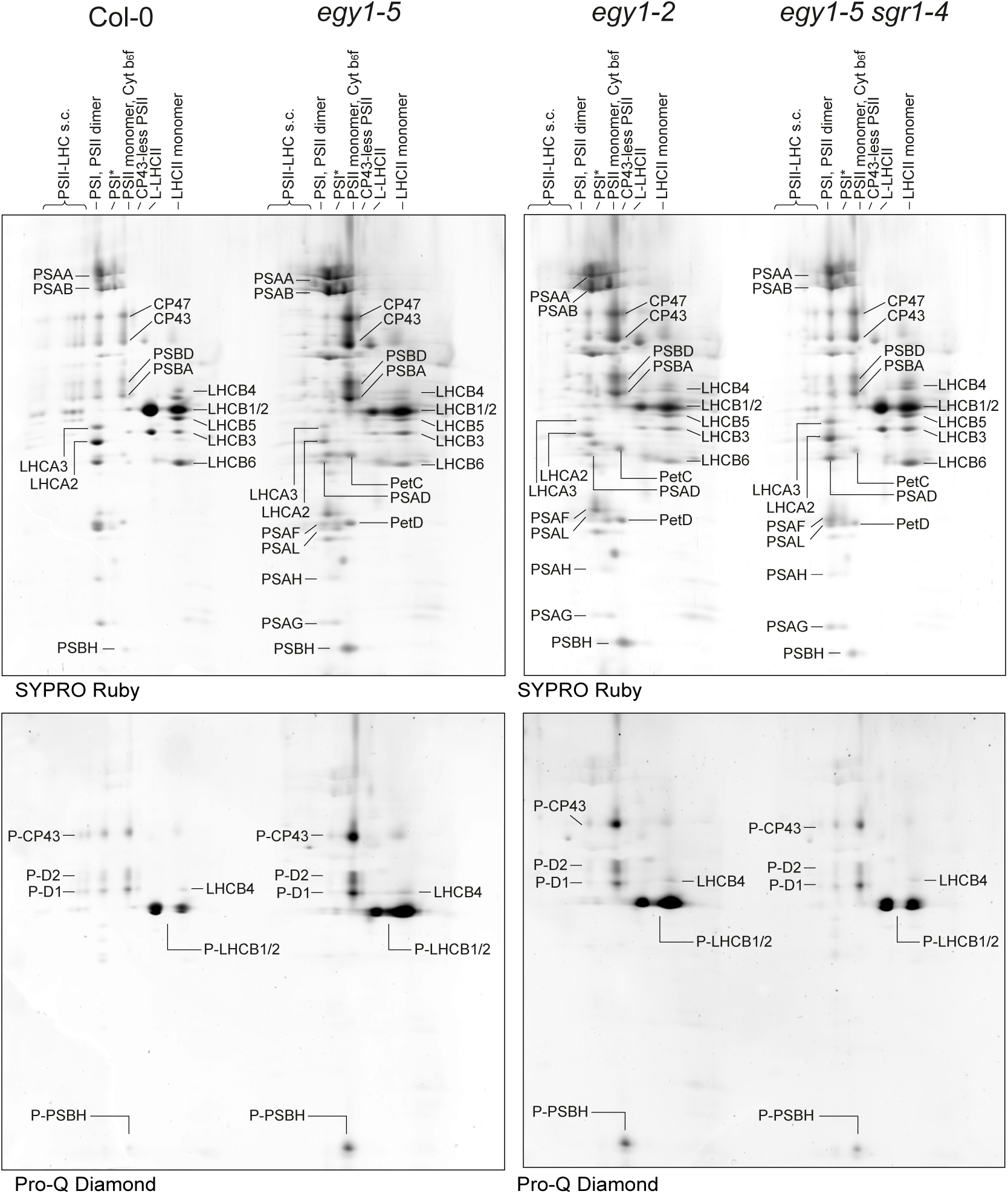
Analysis by 2D-lpBN/SDS-PAGE of the subunit composition of the protein complexes from Col-0 vs *egy1-5* (left panels) and *egy1-2* vs *egy1-5 sgr1-4* (right panels). Gels were stained first with ProQ Diamond for phosphoprotein detection (bottom panels) and subsequently with Sypro to detect all proteins (top panels). The major protein complexes from lpBN-PAGE first dimension are indicated. The protein spots identities are indicated according to (Trotta *et al*., 2025).

**Fig. S7.**
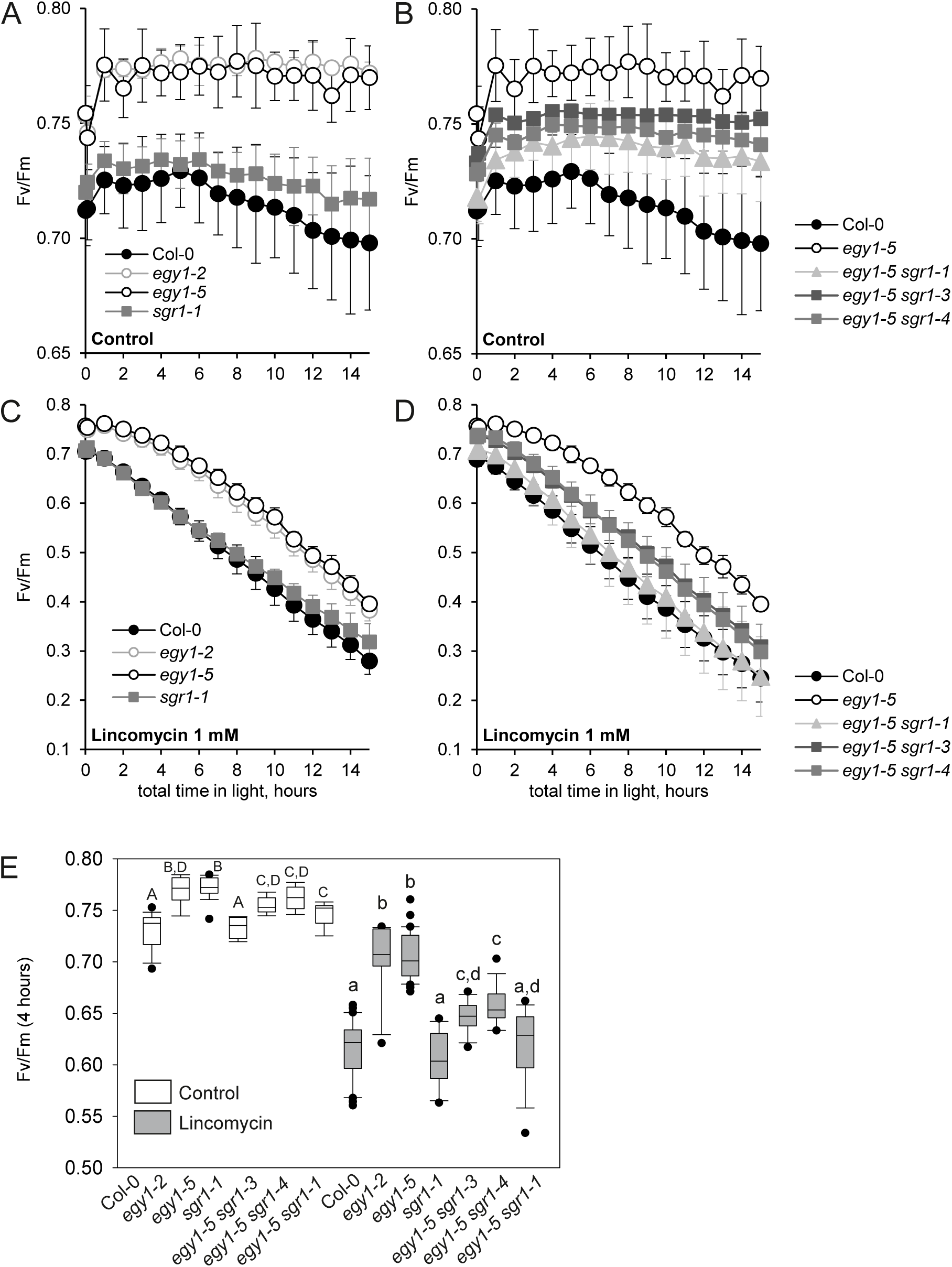
The response to lincomycin. (A-D) Fv/Fm measured in leaf discs of Col-0, *egy1-2*, *egy1-5*, *egy1-5 sgr1*, *egy1-5 sgr1-3, egy1-5 sgr1-4* and *sgr1* exposed to repeated cycles of blue light (448nm, 80 µmol m-2 s-1) under control conditions (A, B) or 1 mM lincomycin treatment (C, D). (E) Data from the 4-hr timepoint was quantified from three biological repeats, N=9 leaf discs for the control treatment and N = 15 leaf discs for the lincomycin treatment. The box plot indicates the median, upper and lower quartile, and dots as outliers. Genotypes with different upper-case letters indicate statistically significant difference in the control, and lower-case letters indicate statistically significant difference in the lincomycin treatment (2-way ANOVA and Tukey’s multiple comparisons test).

**Fig. S8.**
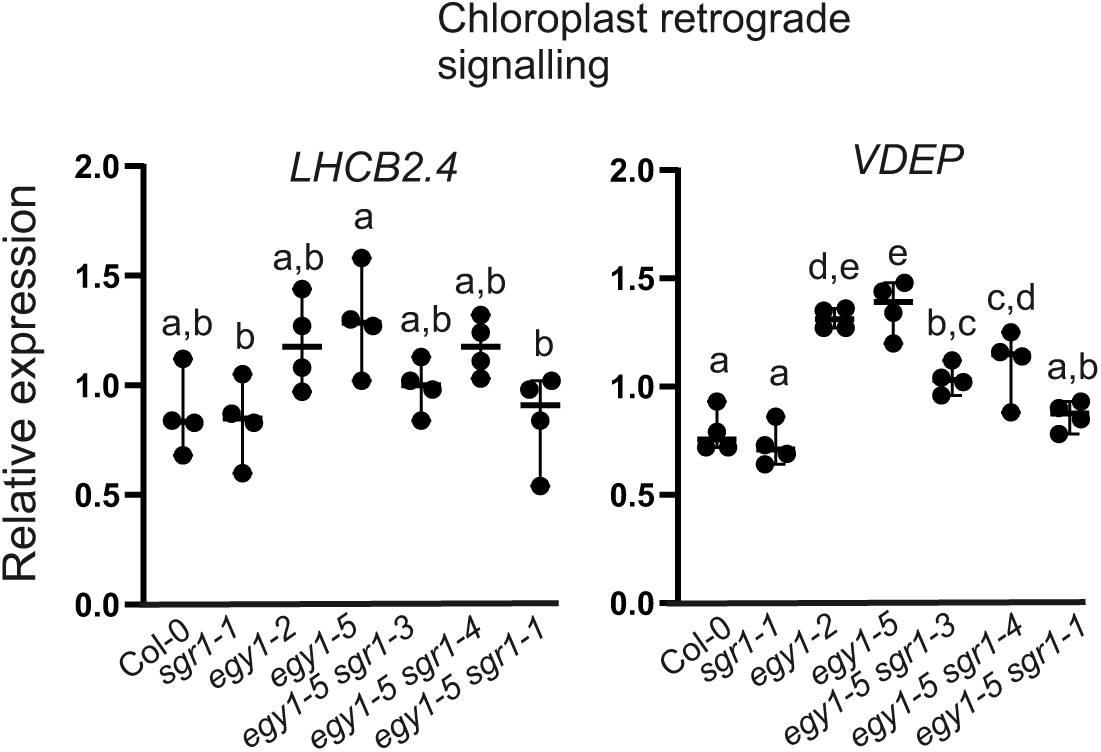
Real-time quantitative PCR (qPCR) was used to assess the relative expression of *LHCB2.4* and *VDEP* and normalized against three reference genes (*PP2AA3*, *TIP41* and *YLS8*). Data represents four biological replicates. Genotypes with different letters are statistically different (1-way ANOVA and Tukey’s multiple comparisons test).

